# The effects of mutation and recombination rate heterogeneity on the inference of demography and the distribution of fitness effects

**DOI:** 10.1101/2023.11.11.566703

**Authors:** Vivak Soni, Susanne P. Pfeifer, Jeffrey D. Jensen

**Author notes:** VS and JDJ conceptualized the project, VS wrote and implemented all code, VS performed the formal analyses with input from SPP and JDJ, and VS, SPP and JDJ wrote the manuscript. This project was funded by National Institutes of Health grant R35GM139383 to JDJ.

## Abstract

Disentangling the effects of demography and selection has remained a focal point of population genetic analysis. Knowledge about mutation and recombination is essential in this endeavour; however, despite clear evidence that both mutation and recombination rates vary across genomes, it is common practice to model both rates as fixed. In this study, we quantify how this unaccounted for rate heterogeneity may impact inference using common approaches for inferring selection (DFE-alpha, Grapes, and polyDFE) and/or demography (fastsimcoal2 and *δaδi*). We demonstrate that, if not properly modelled, this heterogeneity can increase uncertainty in the estimation of demographic and selective parameters and in some scenarios may result in mis-leading inference. These results highlight the importance of quantifying the fundamental evolutionary parameters of mutation and recombination prior to utilizing population genomic data to quantify the effects of genetic drift (*i.e*., as modulated by demographic history) and selection; or, at the least, that the effects of uncertainty in these parameters can and should be directly modelled in downstream inference.

**Significance Statement:** Despite evidence from numerous species that mutation and recombination rates vary along the genome, both rates tend to be modelled as fixed when performing population genetic inference. The impact of failing to account for this rate heterogeneity on the estimation of demographic and selective parameters has yet to be well quantified; thus, we here study this effect by comparing inference under both fixed and variable rate scenarios. Our results demonstrate that unaccounted for rate heterogeneity can increase uncertainty and lead to mis-inference in certain scenarios. We highlight the importance of utilizing mutation and recombination rate maps where possible, and of modelling the uncertainty underlying these estimates.

## Introduction

Quantifying the relative importance of selective and neutral processes in shaping genetic variation has long been a focal point of population genetics. Over the past five decades, the most commonly used null against which hypotheses are compared has remained The Neutral and Nearly Neutral Theories of Molecular Evolution, which together assert that the great majority of newly arising mutations are either neutral or so weakly deleterious that their fixation probabilities will be governed largely by genetic drift, or sufficiently deleterious such that their population-level dynamics will be governed primarily by purifying selection (Kimura 1968, 1983; Ohta 1974). The Neutral and Nearly Neutral Theories also posit a fraction of newly arising beneficial mutations, but suggest that this fraction is small relative to the neutral and deleterious classes of mutations. This general view of an underlying bimodal shape of the distribution of fitness effects (DFE), while controversial at the time, has since been overwhelmingly validated (Jensen et al. 2019; and see reviews of Eyre-Walker and Keightley 2007; Bank et al. 2014).

Subsequent to Kimura’s initial work came the realization that the substantial fraction of sites experiencing selection will also have an important impact on linked sites (see review of Charlesworth and Jensen 2021). Specifically, background selection (BGS) effects may arise at sites linked to targets of purifying selection (Charlesworth et al. 1993), and selective sweep effects may arise at sites linked to targets of positive selection (Maynard Smith and Haigh 1974) – with both effects being types of genetic hitchhiking. As such, the expectations of the Neutral Theory framework would suggest widespread background selection and rare, episodic selective sweeps – expectations entirely consistent with current data (Jensen et al. 2019).

Given this, Johri et al. (2022a, 2022b) recently proposed the necessity of an evolutionary baseline model that accounts for these and other commonly occurring evolutionary processes – including purifying and background selection effects, genetic drift as partially modulated by population history, as well as mutation and recombination rate heterogeneity – in order to more accurately scan for the action of rare processes (*e.g.,* positive and balancing selection). This is particularly important as multiple processes have overlapping predictions in terms of expected levels and patterns of variation, such that, for example, selective effects may often be confused with, and confounded by, demographic effects (*e.g.,* Charlesworth et al. 1993; Charlesworth 1996; Jensen et al. 2005; Kaiser and Charlesworth 2009; O’Fallon et al. 2010; Charlesworth 2013; Nicolaisen and Desai 2013; Zeng 2013; Ewing and Jensen 2016; Johri et al. 2021; Soni et al. 2023).

This circular problem of demography potentially biasing DFE inference and selection potentially biasing demographic inference is relevant to commonly used ’two-step approaches’ for inferring population history and the DFE. In general terms, these approaches seek to infer demography utilizing putatively neutral exonic sites in the genome to infer parameters of the underlying demographic history. Conditional on the inferred demography, the site frequency spectra (SFS) at putatively functional sites is then used to estimate the DFE (*e.g.,* Williamson et al. 2005; Keightley and Eyre-Walker 2007; Li et al. 2012; and see review of Johri et al. 2022a). The benefit of using putatively neutral sites in exonic regions is that they are more likely to share similar mutation and recombination rate environments as the adjacent directly selected sites under investigation. However, one critical assumption with such approaches is that all sites are independent and unlinked, such that genetic hitchhiking effects can be neglected. However, as the effects of selection at linked sites have been shown to be substantial in many organisms (Cutter and Payseur 2013; Charlesworth and Jensen 2021) – primarily in the form of background selection – this framework may be problematic. Indeed, considerable evidence has accumulated demonstrating that this neglect of hitchhiking effects may bias the inference of demography, which will in turn bias the downstream inference of the DFE (*e.g.,* Johri et al. 2021). More specifically, BGS effects may result in a skew in the SFS, with the extent of this skew depending on the underlying DFE. This skew may then by mis-inferred as population size change using neutral demographic estimators, which may in turn result in a mis-inference of the DFE, as DFE inference is predicated upon the demographic inference (Johri et al. 2021; Charlesworth and Jensen 2021).

An alternative to two-step inference approaches involves jointly and simultaneously estimating parameters of demography and selection, without defining an *a priori* neutral / unlinked class of sites (*e.g.,* Johri et al. 2020). Such approaches can directly model the effects of selection at linked sites, whilst also accounting for locus-specific recombination and mutation rates which can act to modify these effects. This class of methods makes use of forward-in-time simulations and approximate Bayesian computation (ABC) to identify a viable model and parameter space that is consistent with observed data. Given this ABC framework and the relaxation of the assumption of independent sites, these methods tend to make use of multiple aspects of the data including the SFS, linkage disequilibrium, and divergence, as opposed to the SFS-only based inference common amongst two-step approaches.

However, the trade-off of simultaneous, joint inference is that large forward-simulation studies are computationally expensive given the parameter and model space that must necessarily be explored. As such, these approaches have thus far been constrained to relatively simple demographic histories (*e.g.,* a single population size change). In addition, whether a two-step or joint inference approach is most appropriate will depend on the underlying details of the study system in question. For instance, the human genome has lengthy intergenic regions separating coding regions, suggesting that the effects of BGS may not be prevalent in certain parts of the genome; as such, carefully constructed two-step approaches may indeed be feasible (Johri et al. 2023). By contrast, the *Drosophila melanogaster* genome contains significantly more functional content, and the effects of selection are likely greater owing to larger effective population sizes, suggesting that a neutral / unlinked class of sites may well not exist – at least in sufficient quantity to estimate a neutral population history – thereby necessitating joint inference (Johri et al. 2020). In organisms that are characterized by almost entirely functional genomes such as many viruses, joint inference will similarly be essential (*e.g.,* Jensen 2021; Morales-Arce et al. 2022; Howell et al. 2023; Terbot et al. 2023).

While these recent developments in joint inference represent important progress in studying demography and selection, less attention has been given to the additional complications of heterogenous mutation and recombination rates when performing evolutionary inference of this sort. Importantly though, it has been demonstrated that demographic inference may be biased by recombination-associated GC-biased gene conversion (Pouyet et al. 2018), whilst the indirect inference of recombination rates from patterns of linkage disequilibrium may be biased by demographic effects (Dapper and Payseur 2018; Samuk and Noor 2022). Furthermore, it is well known that the impact of both background selection and selective sweeps is expected to be stronger in regions of low recombination (Begun and Aquadro 1992; Charlesworth et al. 1993; Wiehe and Stephan 1993; Hudson and Kaplan 1995) – a pattern observed across a wide variety of taxa (see review of Smukowski and Noor 2011). Thus, these results combined with the fact that recombination rates tend to vary across genomes (see review of Stapley et al. 2017), suggests that neglecting this underlying heterogeneity may be problematic when performing evolutionary inference.

Similarly, the rate at which mutations appear throughout the genome varies considerably at every scale examined, from individual sites in a genome to differences observed between individuals, populations, and species (see reviews of Baer et al. 2007; Lynch 2010; Hodgkinson and Eyre-Walker 2011; Pfeifer 2020). Thereby, numerous factors can affect the mutation rate, including: local nucleotide composition (such as GC-content; Hwang and Green 2004), genomic factors (such as chromatin accessibility, replication timing, and recombination; Agarwal and Przeworski 2019), as well as life history traits (see discussion in Tran and Pfeifer 2018). Methods that employ different site categories for demographic and/or DFE inference (Piganeau and Eyre-Walker 2003; Eyre-Walker and Keightley 2009; Gutenkunst et al. 2009; Galtier 2016; Tataru et al. 2017; Tataru and Bataillon 2020) may thus be biased by neglecting this heterogeneity, as underlying rates may differ systematically between different site classes.

Here forward-in-time simulations were run in SLiM (Haller and Messer 2019) to investigate how mutation and recombination rate heterogeneity may impact common demographic and DFE inference procedures (using the software packages *δaδi* [Gutenkunst et al. 2009] and fastsimcoal2 [Excoffier et al. 2013], as well as DFE-alpha [Keightley and Eyre-Walker 2007], Grapes [Galtier 2016], and poly-DFE [Tataru and Bataillon 2020], respectively). We firstly quantify performance under constant-rate assumptions for varying demographic histories and DFE shapes, and then consider changes in performance when only the mutation or recombination rate is variable, as well as when both rates are variable. We show that mutation and recombination rate heterogeneity has important consequences for demographic and DFE inference, and thus should be accounted for in baseline model construction.

## Results and Discussion

### Demographic inference of a neutrally evolving population

Population genetic data was generated and analyzed using forward-in-time simulations in SLiM (Haller and Messer 2019), with the coalescent approach fastsimcoal2 (Excoffier et al. 2013), and with the diffusion approximation approach implemented in *δaδi* (Gutenkunst et al. 2009). As well as being widely used for demographic inference, fastsimcoal2 and *δaδi* use differing approaches that are both implemented in a likelihood framework, and are therefore a useful comparison. Initially, a neutrally evolving equilibrium population was simulated under four different rate-based scenarios: (1) fixed population-scaled recombination rate (*ρ*) and population-scaled mutation rate (*θ*), (2) fixed *ρ* and variable *θ*, (3) variable *ρ* and fixed *θ*, as well as (4) variable *ρ* and variable *θ*. Fixed rates were taken from the *D. melanogaster* genome-wide averages (Comeron et al. 2012 and Keightley et al. 2014 for *ρ* = 2*Nr* and *θ* = 2*Nμ*, respectively, where *N* is the ancestral population size [*N_ancestral_*], *μ* is the per-site per-generation mutation rate, and *r* is the per-site per-generation recombination rate), and variable rates were drawn from a uniform distribution, such that the mean rate across each simulation replicate was equal to the fixed rate, to enable fair comparisons (see the Materials and Methods section for further details). 100 replicates were simulated for each rate regime, and then combined into 10 SFS, each consisting of 10 combined replicates (as per the approach of Excoffier et al. 2013). Demographic inference was performed on each of these spectra using fastsimcoal2 (Excoffier et al. 2013) and *δaδi* (Gutenkunst et al. 2009). Simulations were also performed under various population size change scenarios whereby the size change was instantaneous and occurred 1*N_ancestral_* generations prior to sampling. Three non-equilibrium scenarios were simulated: (1) an instantaneous 100% expansion, (2) an instantaneous 50% contraction, and (3) a more severe instantaneous 99% contraction (see Supplementary Figure S1 for schematics of the demographic histories simulated). fastsimcoal2 infers the population size at time of sampling (*N_current_*), as well as – when a population size change model is used – the ancestral population size (*N_ancestral_*) and the time of population size change (*τ_ancestral_*). By contrast, *δaδi* infers the population size at time of sampling relative to the ancestral population size (*Nu_opt_*), and the time of population size change relative to the ancestral population size (*τ_opt_*). All demographic inference results were scaled relative to the true parameter values (for details see Methods).

Both methods performed well when inferring the population size at time of sampling for the equilibrium population scenario, as well as for the ancestral population size (in the case of fastsimcoal2) and time of size change (Figure 1), consistent with previous studies (Excoffier et al. 2013; Johri et al. 2021), regardless of whether mutation and/or recombination rates were fixed or variable (and see Supplementary Figures S2-S5 for summary statistics corresponding to the simulation results in Figure 1). However, it is notable that the variance increased greatly for estimates of *N_current_* (the population size at time of sampling) and *N_ancestral_* (the initial population size) with fastsimcoal2 for the severe population contraction scenario. Variable recombination rates had little effect on nucleotide diversity or the number of segregating sites under this neutral model, while variable mutation rates increased the variance of the underlying statistics (Supplementary Figures S2-S5).

**Figure 1:**
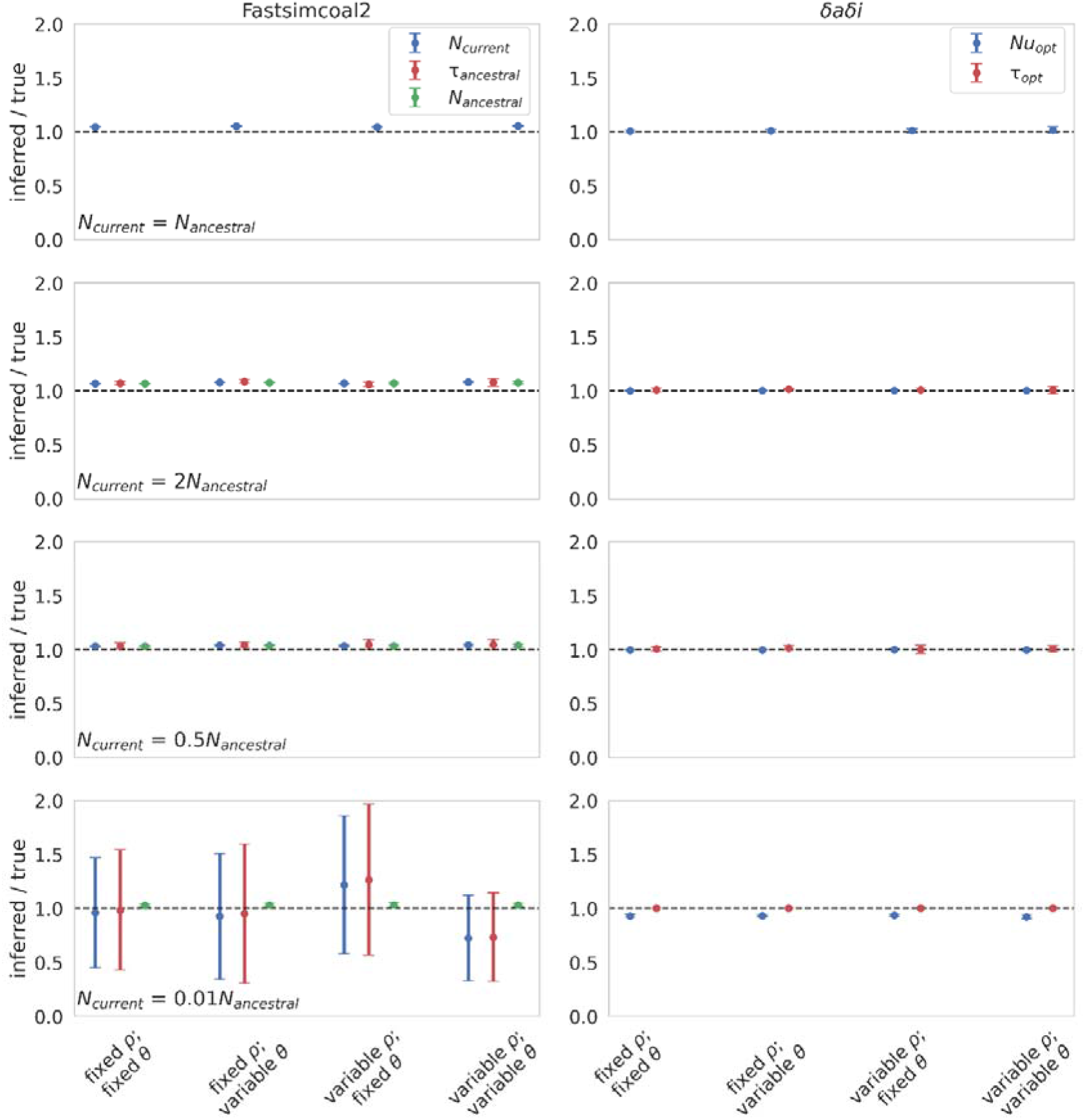
Demographic inference results for a single neutrally evolving population under fixed and variable recombination and mutation rates. Each row depicts a different demographic history: equilibrium population; instantaneous 100% expansion; instantaneous 50% contraction; and instantaneous 99% contraction. Population size change occurred 1*N_current_* generations before sampling, where *N_current_* is the population size at time of sampling. All inference values are scaled by the true value. As such, a value of 1 on the Y-axis indicates that the inferred parameter value matches the true parameter value (as shown by the black dashed line). fastsimcoal2 infers three parameters: *N_current_*; *N_ancestral_* (the initial population size); and *τ_ancestral_* (the time of population size change in *N_current_* generations). Only *N_current_* was inferred for the equilibrium population scenario. *δaδi* infers two parameters: *Nu_opt_* (the population size at time of sampling relative to the initial population size); and *τ_opt_* (the time of population size change relative to the initial population size). Data points represent the mean inference value across 10 replicates, whilst error bars represent the standard deviation. *θ = 2Nμ* and *ρ = 2Nr,* where *N* is the ancestral population size (*N_ancestral_*), *μ* is the per-site per-generation mutation rate, and *r* is the per-site per-generation recombination rate. Variable rates are drawn from a uniform distribution, such that the mean rate across each simulation replicate was equal to the fixed rate, to enable fair comparisons (see the Materials and Methods section for further details, and Supplementary File S1 for inference means, standard deviations, and quartile values).

These results were as expected, given that the population was neutrally evolving. Specifically, the coalescent approach of fastsimcoal2 (Excoffier et al. 2013) should be robust to mutation and recombination rate variation in the absence of selection, because the rate of coalescence should remain unaffected (Wakeley 2009), whilst estimators such as *δaδi* (which uses a composite likelihood approach to compare the estimated and observed data; Gutenkunst et al. 2009) have been shown to be consistent for selectively neutral data under a wide range of population genetic scenarios (Wiuf 2006). However, a more realistic model would include the effects of both purifying and background selection, and thus, we next examined these models in the presence of a DFE at functional sites.

### Demographic inference in an equilibrium population, in the presence of purifying and background selection

To incorporate the effects of purifying and background selection, functional elements were simulated, each separated by intergenic regions (see Materials and Methods and Supplementary Figure S6 for simulation details and demographic history schematics, respectively), with intergenic and intronic regions evolving neutrally. Within exonic regions, mutations were draw from a discrete DFE, and six separate DFEs were evaluated (following Johri et al. 2020) as depicted in Figure 2. The six DFEs included: (1) an excess of weakly deleterious mutations; (2) an excess of moderately deleterious mutations; (3) an excess of strongly deleterious mutations; (4) an equal distribution of selection coefficients across mutations; as well as (5) and (6) bimodal distributions, as are most commonly inferred in empirical / experimental DFE estimates for newly arising mutations. These enabled us to evaluate demographic inference under multiple purifying and background selection regimes.

**Figure 2:**
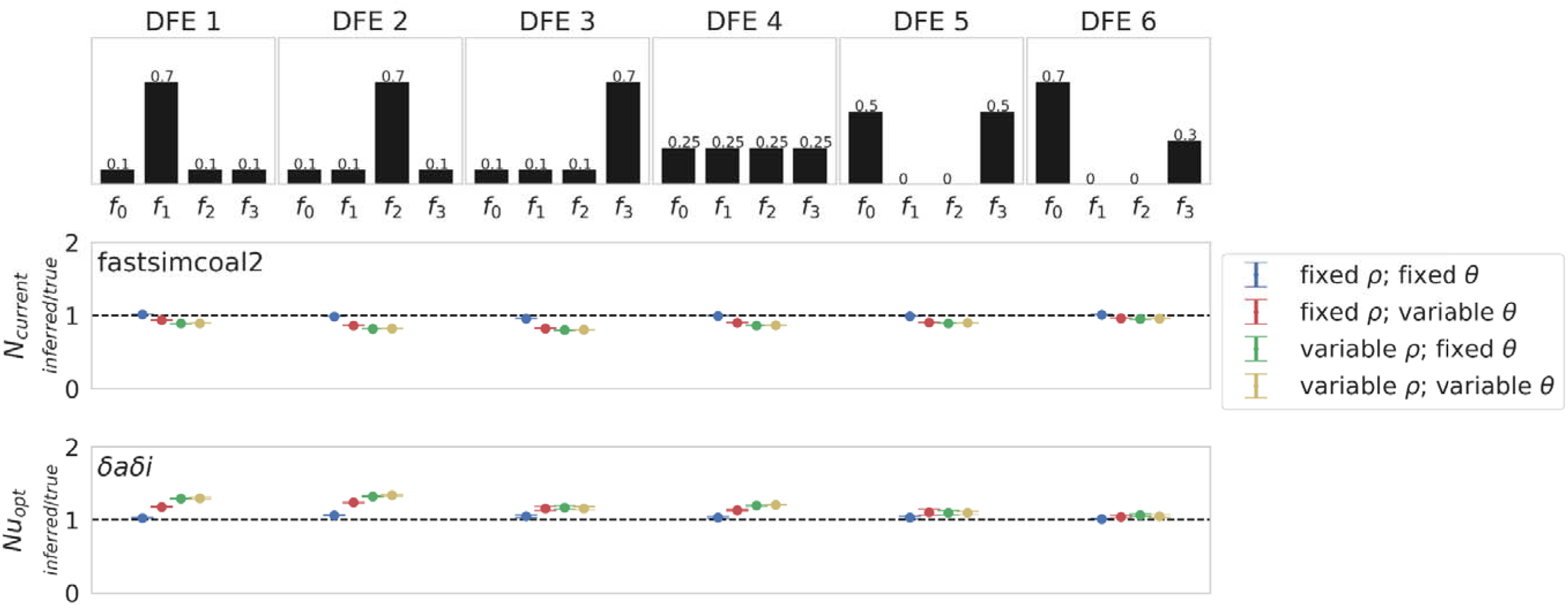
Demographic inference results for a single equilibrium population under multiple distributions of fitness effects (DFEs), with exonic regions masked. Top row depicts the six discrete DFEs that were simulated. Exonic mutations were drawn from a DFE comprised of four fixed classes (Johri et al. 2020), whose frequencies were denoted by *f_i_: f_0_* with 0 ≤ *2N_ancestral_ s* < 1 (*i.e.*, effectively neutral mutations), *f_1_* with 1 ≤ 2*N_ancestral_ s* < 10 (*i.e.*, weakly deleterious mutations), *f_2_* with 10 ≤ 2*N_ancestral_ s* < 100 (*i.e.*, moderately deleterious mutations), and *f_3_* with 100 ≤ 2*N_ancestral_ s* (*i.e.*, strongly deleterious mutations), where *N_ancestral_* was the initial population size and *s* was the reduction in fitness of the mutant homozygote relative to wild-type. Middle and bottom rows show inference results for fastsimcoal2 and *δaδI,* respectively. All inference values are scaled by the true value. As such, a value of 1 on the Y-axis indicates that the inferred parameter value matches the true parameter value (as shown by the black dashed line). For the equilibrium population model, fastsimcoal2 infers the *N_current_* parameter, which is the population size at time of sampling, whilst *δaδi* infers *Nu_opt_*, which is the population size at time of sampling relative to the initial population size. Data points represent the mean inference value across 10 replicates, whilst error bars represent the standard deviation. *θ = 2Nμ* and *ρ = 2Nr,* where *N* is the ancestral population size (*N_ancestral_*), *μ* is the per-site per-generation mutation rate, and *r* is the per-site per-generation recombination rate. Variable rates were drawn from a uniform distribution, such that the mean rate across each simulation replicate was equal to the fixed rate, to enable fair comparisons (see the Materials and Methods section for further details, and Supplementary File S1 for inference means, standard deviations, and quartile values).

Demographic inference was once more performed using fastsimcoal2 (Excoffier et al. 2013) and *δaδi* (Gutenkunst et al. 2009). Inference was performed on both the entire simulated region (see Supplementary Figure S7) as well as separately with exonic regions masked (Figure 2), as is standard practice (Gutenkunst et al. 2009; Excoffier et al. 2013). By masking exonic regions, the effects of purifying selection on inference should be minimized (Charlesworth et al. 1993; Charlesworth 1996; Kaiser and Charlesworth 2009; O’Fallon et al. 2010; Charlesworth 2013; Nicolaisen and Desai 2013). However, background selection can still impact linked sites, in a manner that will depend on the strength of selection and the rate of recombination (Charlesworth et al. 1993; Charlesworth 1996; and see review of Charlesworth and Jensen 2021).

As shown in Figure 2, fastsimcoal2 underestimated *N_current_* when there was an excess of moderately or strongly deleterious mutations (DFEs 2 and 3), and mutation and/or recombination rates were variable, likely due to the increased variance in the genealogies being interpreted as stronger genetic drift effects and therefore a reduced population size. This directionality of bias was further supported by the greater underestimation of *N_current_* when functional sites were not masked (Supplementary Figure S7). Inference with *δaδi* generally followed an opposite pattern (Figure 2). When mutation and recombination rates were fixed, a small overestimate of *Nu_opt_* was observed when there was an excess of moderately or strongly deleterious mutations (DFEs 2 and 3). Variable rates increased the overestimate, most notably in DFEs 1 and 2 (which exhibited an excess of weakly deleterious or moderately deleterious mutations), likely owing to the resulting left-skew in the SFS (*e.g.,* a negative value of Tajima’s *D* [Tajima 1989]; Supplementary Figure S8, and see Supplementary Figure S9 for summary statistics when exonic regions are unmasked). This excess of rare alleles segregating under these DFEs was further exacerbated in low recombination regions where the effects of BGS have a longer range (Charlesworth et al. 1993; Campos and Charlesworth 2019; and see review of Charlesworth and Jensen 2021). As *δaδi* utilizes the shape of the SFS, this left skew was likely interpreted as population growth (*i.e.,* a larger *N_current_*).

### Demographic inference under population size change, in the presence of purifying and background selection

Further simulations incorporated three instantaneous population size change scenarios (100% expansion, 50% contraction, and 99% contraction – see Supplementary Figure S1) with the six DFEs described above (see Figure 2).

Both fastsimcoal2 and *δaδi* estimated all parameters well for the 100% expansion and 50% contraction scenarios when recombination and mutation rates were constant across the genome (*i.e.,* the scenario for which they were designed; see Figure 3 for population expansion, and Supplementary Figures S10-11 for 50% and 99% contractions, respectively); by comparison, the variance on fastsimcoal2 inference of *N_current_* and *τ_ancestral_* greatly increased under the 99% contraction scenario, and *δaδi* overestimated both *Nu_opt_* and *τ_ancestral_*. Under a model of population expansion, all three parameters were underestimated by fastsimcoal2 when rates were variable (Figure 3; and see Supplementary Figure S12 for summary statistics), suggesting a more recent expansion, particularly when recombination rates were variable. By contrast, *δaδi* tended to overestimate *N_current_*. These results likely owe to the fact that population expansion is expected to skew the SFS towards rare alleles (as supported by the reduction in Tajima’s *D* [Tajima 1989]; Supplementary Figure S12) – a pattern that may be further exacerbated by BGS in the presence of weakly deleterious mutations. Results appear to support this thesis, with both methods underestimating *τ_ancestral_*, most notably in DFE2, in which there existed an excess of weakly deleterious mutations. These factors together generated a considerable underestimate of the time since the population size change in DFE2. For DFEs in which no weakly deleterious mutations were segregating (DFEs 5 and 6), the effect was less pronounced. Indeed, *δaδi* inferred *τ_opt_* well for all other DFEs. As with the equilibrium population simulations (Figure 2), the population size at time of sampling was underestimated by fastsimcoal2, and overestimated by *δaδi* when rates were variable.

**Figure 3:**
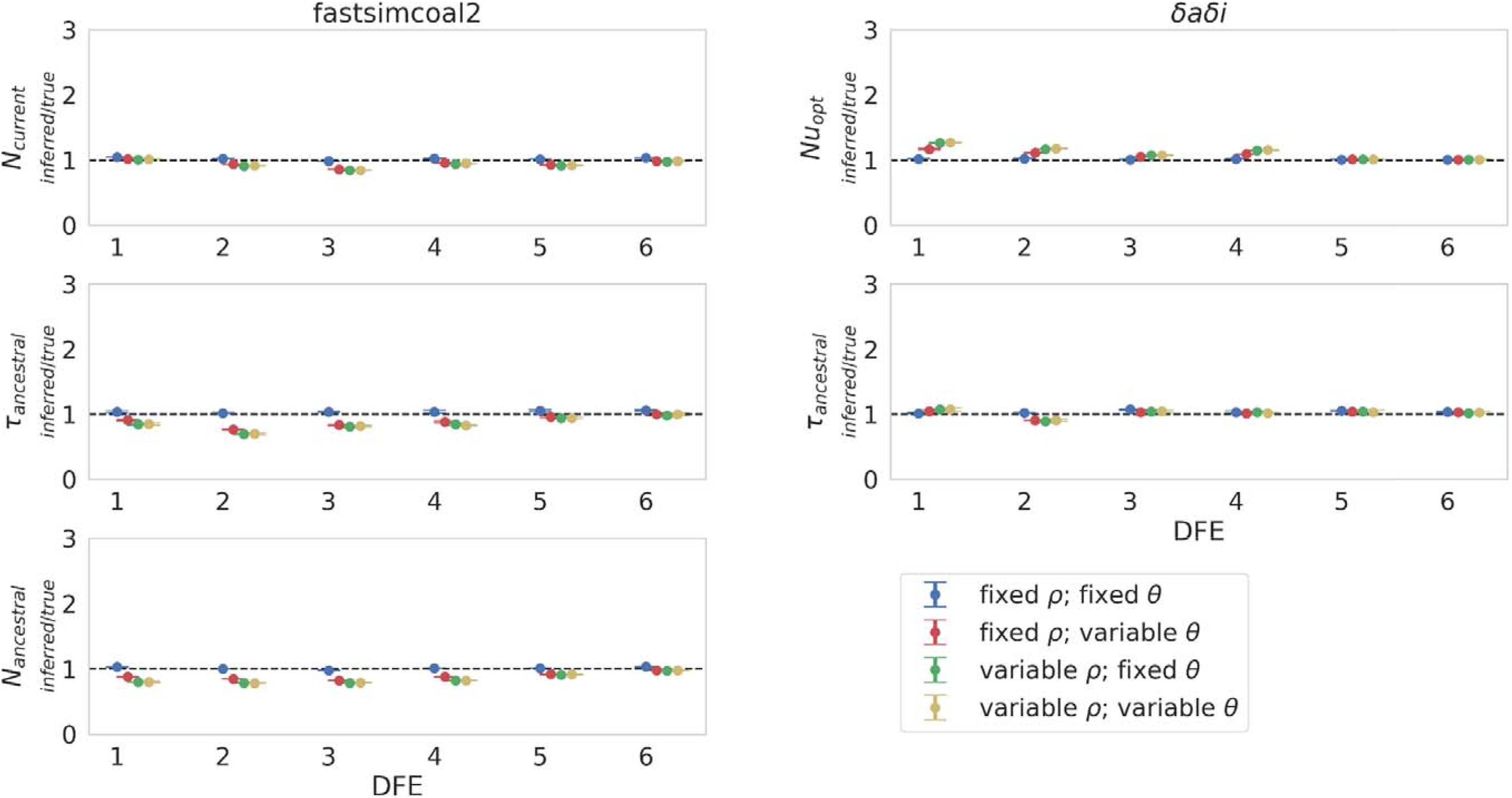
Demographic inference results for a population expansion (such that *N_current_* = 2*N_ancestral_*) under fixed and variable recombination and mutation rates, across the six simulated distributions of fitness effects (DFEs), with exonic regions masked. Population size change occurred 1*N_current_* generations before sampling, where *N_current_* is the population size at time of sampling. Exonic mutations were drawn from a DFE comprised of four fixed classes (Johri et al. 2020), whose frequencies were denoted by *f_i_: f_0_* with 0 ≤ 2*N_ancestral_ s* < 1 (*i.e.*, effectively neutral mutations), *f_1_* with 1 ≤ 2*N_ancestral_ s* < 10 (*i.e.*, weakly deleterious mutations), *f_2_* with 10 ≤ 2*N_ancestral_ s* < 100 (*i.e.*, moderately deleterious mutations), and *f_3_* with 100 ≤ 2*N_ancestral_ s* (*i.e.*, strongly deleterious mutations), where *N_ancestral_*was the initial population size and *s* was the reduction in fitness of the mutant homozygote relative to wild-type. fastsimcoal2 infers three parameters: *N_current_*; *N_ancestral_*; and *τ_ancestral_* (the time of population size change in *N_current_* generations). *δaδI* infers two parameters: *Nu_opt_* (the population size at time of sampling relative to the initial population size); and *τ_opt_* (the time of population size change relative to the initial population size). Data points represent the mean inference value across 10 replicates, whilst error bars represent the standard deviation. *θ = 2Nμ* and *ρ = 2Nr,* where *N* is the ancestral population size (*N_ancestral_*), *μ* is the per-site per-generation mutation rate, and *r* is the per-site per-generation recombination rate. Variable rates are drawn from a uniform distribution, such that the mean rate across each simulation replicate was equal to the fixed rate, to enable fair comparisons (see the Materials and Methods section for further details, and Supplementary File S1 for inference means, standard deviations, and quartile values).

With population contraction, an increase in the variance of the underlying genealogies is expected (Hein et al. 2005), and combined with background selection, this resulted in an overestimate of the time of population size change. When rates were fixed this overestimate was slight with both methods for a 50% population contraction (Supplementary Figure S10; see Supplementary Figure S13 for summary statistics), though variable rates (particularly recombination rates) increased the overestimate, most notably in DFEs in which weakly or moderately deleterious mutations were segregating. The dynamics of inference of the population size at time of sampling were similar to the previously discussed scenarios, where **N_current_** was underestimated by fastsimcoal2, whilst *Nu_opt_* was overestimated by *δaδi* when rates were variable. Under the more severe 99% population contraction (Supplementary Figure S11; see Supplementary Figure S14 for summary statistics), the variance in *N_current_*and *τ_ancestral_* with fastsimcoal2 greatly increased, whilst *δaδi* tended to overestimate both *Nu_opt_* and *τ_ancestral_*. However, the two methods accurately inferred a general model consisting of a recent and severe population contraction. The SFS for this demographic history is skewed severely towards rare alleles (see Supplementary Figure S14 for Tajima’s *D* estimates), though the variance on this skew is extremely high regardless of whether rates are fixed or variable, and this variance is likely a driving factor for generating mis-inference (for inference results and summary statistics with no exonic masking, see Supplementary Figures S15-20).

### DFE inference

DFE inference was performed using two methods that utilize the widely used approach of Keightley and Eyre-Walker (2007): DFE-alpha (Keightley and Eyre-Walker 2007) and Grapes (Galtier 2016); and the more recently proposed poly-DFE approach (Tataru and Bataillon 2020) was evaluated for comparison. Inference was performed on the six simulated DFEs shown in Figure 2, with the methods inferring a gamma-distributed DFE of new mutations and then calculating the proportion of mutations in each of four fixed DFE classes (see Materials and Methods).

Both DFE-alpha and Grapes performed well when mutation and recombination rates were fixed, particularly for DFEs 1-3, which are nearer to a gamma distribution than DFEs 4-6 (Figure 4; see Supplementary Figure S21 for Grapes inference results). However, when recombination and/or mutation rates were variable, the inferred DFEs were largely neutral (*i.e.,* biased towards *f_0_* mutations), with a small proportion of weakly deleterious mutations. This pattern was consistent across the six simulated DFEs, and across population histories (although DFE-alpha inferred a completely neutral DFE for population contraction scenarios). These results suggest that both methods struggle to find the maximum likelihood value of the mean parameter under variable rate scenarios. When rates are fixed, inference was modestly affected by moderate demographic size changes (Supplementary Figures S22-S25). Under a 99% contraction (Supplementary Figures S26-S27), a bias towards strongly deleterious mutations was observed with DFE-alpha, whilst Grapes appeared more robust to such a population size change. This difference in inference may reflect differences in how the methods account for the effects of demography; while Grapes re-weights the SFS (Eyre-Walker et al. 2006; Galtier 2009), DFE-alpha fits DFE models to the ratio of the SFS at selected vs. putatively neutral sites (Eyre-Walker and Keightley 2007). Indeed, approaches which assume that demography impacts neutral and selected polymorphisms to the same extent have been shown to be biased under severe population size changes (Eyre-Walker et al. 2006). The likelihoods of the neutral and *GammaZero* models implemented in Grapes were compared (see Supplementary File S2 for likelihood values for neutral vs *GammaZero* models), revealing a largely consistent pattern in which the *GammaZero* model was the more likely for DFEs 1-4 when rates were fixed, but the model was rejected for DFEs 5 and 6. Under variable rates, the *GammaZero* model was always rejected, across all population histories.

**Figure 4:**
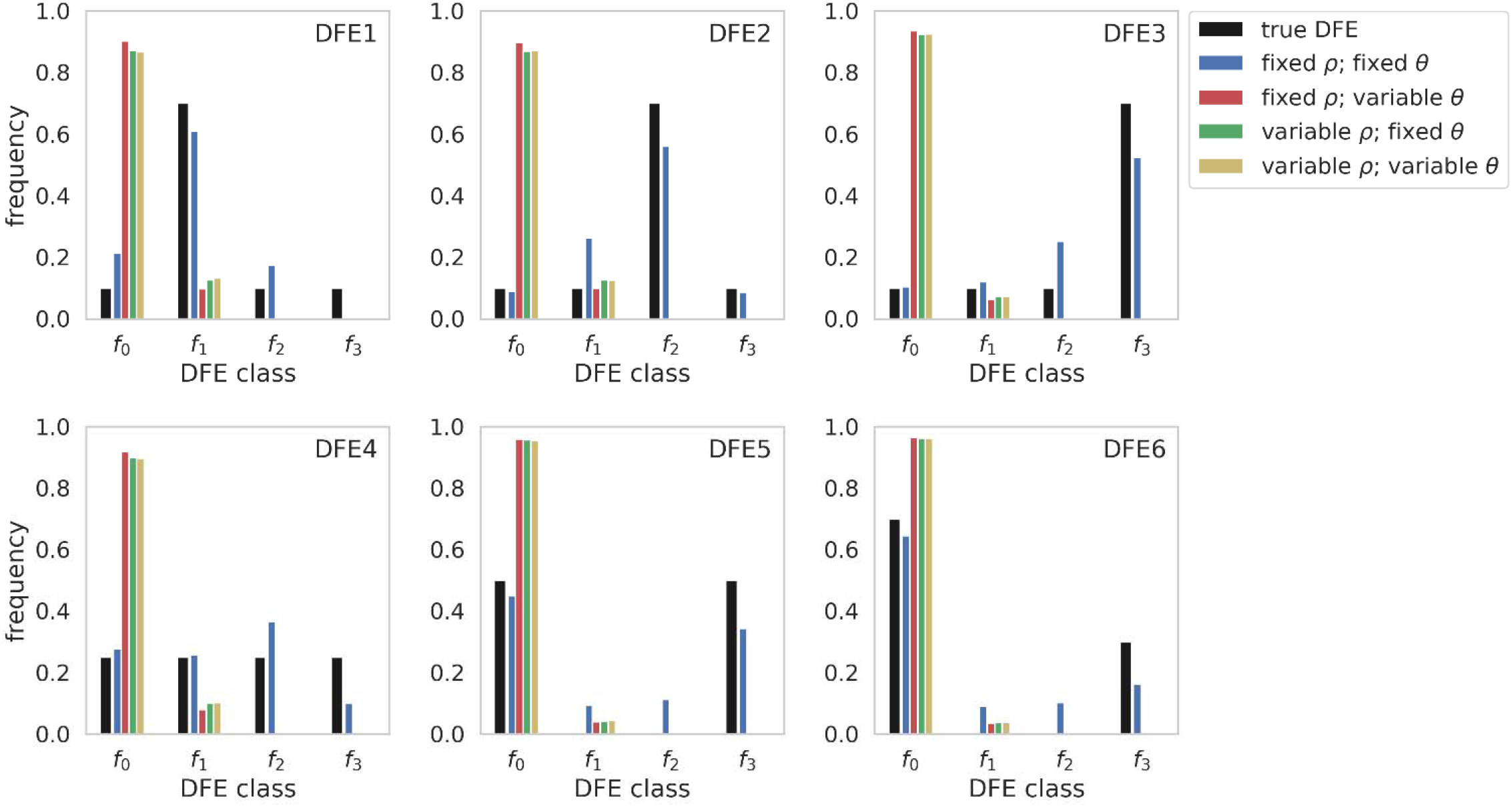
Comparison of true distributions of fitness effects (DFEs) and DFE inference using DFE-alpha, when recombination and mutation rates were fixed and/or variable, under the six DFE models, for a constant-size population. Exonic mutations were drawn from a DFE comprised of four fixed classes (Johri et al. 2020), whose frequencies were denoted by *f_i_: f_0_* with 0 ≤ 2*N_ancestral_ s* < 1 (*i.e.*, effectively neutral mutations), *f_1_* with 1 ≤ 2*N_ancestral_ s* < 10 (*i.e.*, weakly deleterious mutations), *f_2_* with 10 ≤ 2*N_ancestral_ s* < 100 (*i.e.*, moderately deleterious mutations), and *f_3_* with 100 ≤ 2*N_ancestral_ s* (*i.e.*, strongly deleterious mutations), where *N_ancestral_* was the initial population size and *s* was the reduction in fitness of the mutant homozygote relative to wild-type. *θ = 2Nμ* and *ρ = 2Nr,* where *N* is the ancestral population size (*N_ancestral_*), *μ* is the per-site per-generation mutation rate, and *r* is the per-site per-generation recombination rate. Variable rates were drawn from a uniform distribution, such that the mean rate across each simulation replicate was equal to the fixed rate, to enable fair comparisons (see the Materials and Methods section for further details).

Because polyDFE (Tataru and Bataillon 2020) allows for mutation rate variation, unlike the above two methods, this approach was applied to our equilibrium population simulations for comparison, where recombination rates were fixed and mutation rates were variable. polyDFE accounts for mutation rate variation by separating each variable rate region into a separate SFS fragment (*i.e.,* each variable rate region is submitted as a separate SFS in the input file). However, inference remained poor, with an excess of effectively neutral and strongly deleterious mutations inferred for all six simulated DFEs (Supplementary Figure S28). This consistency of results amongst the three DFE estimators likely owes to their shared inability to account for the effects of BGS; specifically, the inability to account for the SFS-skewing generated by weakly and moderately deleterious mutations. Furthermore, the variance of this effect is inflated when mutation rates are variable.

### Concluding thoughts

Distinguishing selective from neutral processes is challenging, because these competing processes may fundamentally result in similar patterns of variation (Barton 2000; Johri et al. 2022b). Perhaps the most common approach to this problem is two-step inference, in which a demographic history is inferred from the SFS at putatively neutral and unlinked sites of the genome, followed by DFE inference at putatively functional sites utilizing the inferred population history. This study has evaluated the effects of recombination and/or mutation rate heterogeneity on such two-step approaches, showing that both processes may significantly bias inference. This result is notable given that many organisms of interest have been observed to have considerable underlying heterogeneity in both mutation and recombination rates across their genomes (see review of Stapley et al. 2017 for recombination rate variation, and reviews of Lynch 2010; Hodgkinson and Eyre-Walker 2011; Pfeifer 2020 for mutation rate variation).

The results presented here demonstrate that the effects of this rate heterogeneity on demographic inference will depend on both the true demographic history as well as the underlying DFE in functional regions. The severity of bias was found to be smaller for the inference of population size, and greater for inferring the timing of population size change. These biases were found to be exacerbated by an excess of weakly deleterious mutations in functional regions, owing to the resulting background selection effects at linked sites. More generally, all neutral demographic approaches are biased by the effects of selection if present in the data, and the reliance on the SFS alone (and the related assumption of unlinked sites) leaves potentially informative patterns (*e.g.,* linkage disequilibrium) unutilized. In the case of fastsimcoal2, the inability to account for the biasing effects of BGS is exacerbated by the increased variance created by rate heterogeneity. In the case of *δaδi,* the assumption of independent sites is being violated to highly varying degrees across datasets under these variable rate regimes.

DFE inference proved more challenging, with variable mutation and recombination rates resulting in the inference of an excess of effectively neutral mutations, regardless of the underlying DFE. Although polyDFE performed slightly better than the other methods with variable mutation rates, the method was still unable to effectively distinguish amongst the six different DFE shapes. These results suggest that the generally neglected fine-scale rate heterogeneity often underlying genomic data may lead to substantial biases when inferring selection particularly.

Taken together, these results highlight the importance of utilizing recombination and mutation rate maps defined for the specific population of interest, prior to performing evolutionary genomic inference of this sort. While this represents a challenge for many non-model organisms for which such fine-scale rates have yet to be well-defined, this suggests such efforts to be necessary steps for accurate downstream inference. Pedigree-based studies are feasible in many organisms, and can be used for both *de novo* mutation rate estimation and recombination rate estimation. Mutation accumulation experiments may also be performed in a smaller subset of lab-tractable organisms (see reviews of Lynch et al. 2016 and Pfeifer 2020 for common mutation rate inference approaches as well as Stumpf and McVean 2003, and Peñalba and Wolf 2020 for common recombination rate inference approaches). Where these approaches are not tractable, the possible inference impact of the uncertainty in underlying mutation and recombination rates can be directly quantified by drawing rates from across the feasible range (*e.g.,* by drawing from appropriate prior distributions, if inference is performed within an approximate Bayesian context (Johri et al. 2022b)).

Importantly however, only by accurately defining these population-specific baseline models of variation consisting of the certain-to-be-operating evolutionary processes of purifying and background selection, population size change and structure, and mutation and recombination rate heterogeneity, may one hope to quantify the relative roles of episodic evolutionary processes such as positive or balancing selection (Johri et al. 2022b). In practice this involves the construction and simulation of baseline models incorporating these common and constantly operating evolutionary processes, in order to determine whether observed empirical data may be fit by these processes alone. The failure to construct a more realistic baseline model of this sort may result in an unjustified dependence on rare or episodic evolutionary processes as explanatory factors.

### Methods and Materials

### Simulations

Forward-in-time simulations were performed using SLiM 3.7 (Haller and Messer 2019), for a single population. Two chromosomal structures were simulated: (1) a strictly neutral model consisting of a single 1Mb region, and (2) a model that incorporates a DFE. For this second model, the numbers and lengths of exons, introns, and intergenic regions were averages estimated from Ensembl’s BDGP6.32 dataset (Adams et al. 2000), obtained from Ensembl release 107 (Cunningham et al. 2022), and were used to simulate a chromosome with *Drosophila melanogaster*-type structure and parameterizations. Specifically, each replicate was made up of 127 genes separated by intergenic regions of size 3,811bp. Each gene contained four exons (of size 588bp) and three introns (of size 563bp), resulting in a gene length of 4,041bp. The total chromosome length was 997,204bp (see Supplementary Figure S6 for a schematic representation of the genic structure).

### Simulating variable recombination and mutation rates

Mutations in intronic and intergenic regions were all effectively neutral, whilst exonic mutations were drawn from a DFE comprised of four fixed classes (following Johri et al. 2020), with frequencies denoted by *f_i_*: *f_0_* with *0 ≤ 2N_ancestral_ s < 1* (*i.e.,* effectively neutral mutations), *f_1_* with *1 ≤ 2N_ancestral_ s < 10* (*i.e.,* weakly deleterious mutations), *f_2_* with *10 ≤ 2N_ancestral_ s < 100* (*i.e.,* moderately deleterious mutations), and *f_3_* with *100 ≤ 2N_ancestral_ s* (*i.e.,* strongly deleterious mutations), where *N_ancestral_*is the initial population size and *s* is the reduction in fitness of the mutant homozygote relative to the wild-type. All mutations were semi-dominant (*h* = 0.5) and, within each bin, *s* was drawn from a uniform distribution. Every third site within exons was simulated to be neutral (*i.e.,* to serve as synonymous sites). Overall, six different discrete DFEs were simulated (for parameterizations, see Figure 2).

The fixed per-site per-generation mutation rate was taken from Keightley et al.’s (2014) estimates from *D. melanogaster* samples. This rate of 2.8 × 10^-9^/site/generation suggests an effective population size of 1.4 million. The fixed recombination rate was taken from Comeron et al.’s (2012) genome-wide average estimate (2.32cM/Mb). Four scenarios were simulated: (1) fixed mutation rate and fixed recombination rate, (2) fixed mutation rate and variable recombination rate, (3) variable mutation rate and fixed recombination rate, as well as (4) variable mutation rate and variable recombination rate. For the fixed rate scenarios, the above rates were constant across the entire simulated chromosome; when variable, each 1kb-region of the simulated chromosome had a different rate drawn from a uniform distribution such that the chromosome-wide average was equal to the fixed rate. For mutation rates, the limits of the uniform distribution were *a*=1e-9 and *b*=6.1e-9 (*i.e.,* the confidence intervals on the Keightley et al. 2014 point-estimate). For recombination rates, the limits of the uniform distribution were *a*=0.0127 and *b*=7.40 (*b* is the maximum rate from Comeron et al. 2012).

### Simulating population size change

Four population histories were simulated: (1) demographic equilibrium, (2) instantaneous 100% population expansion, (3) instantaneous 50% population contraction, and (4) instantaneous 99% population contraction. In each case, 100 chromosomes were sampled after a 16*N_ancestral_* generation burn-in, followed by the size change, followed by 1*N_current_*generations. 100 simulation replicates were generated for each scenario, and all parameters were scaled down 200-fold in order to reduce runtimes, resulting in an ancestral population size of *N_ancestral_* = 7,000. This necessarily includes the scaling up of all parameters that are scaled down by *N* (including mutation rates, recombination rates, and selection coefficients; see Hill and Robertson 1966).

### Demographic inference with fastsimcoal2

Demographic inference with fastsimcoal2 (version fsc27) was performed on all single nucleotide polymorphism (SNP) data, and separately with exonic SNPs masked. A SFS was constructed for each batch of 10 simulation replicates, giving a total of 10 spectra, each covering a 10Mb (for neutral simulations) or 9.97Mb (for DFE simulations) region. fastsimcoal2 was used to fit each SFS to a demographic model. For neutral demographic simulations, SFS were fitted to the equilibrium model, which estimates a single population size parameter (*N_current_*). For instantaneous population size change, simulated spectra were fitted to the instantaneous size change (growth/decline) model, which fits three parameters: ancestral population size (*N_ancestral_*), current population size (*N_current_*), and time of change (*τ*). The parameter search ranges for *N_ancestral_* and *N_current_* were specified to be uniformly distributed between 10 and 100,000 individuals, whilst the range for *τ* was specified to be uniform between 10 and 10,000 generations. For each simulation replicate inference was conducted 100 times, with 100 optimization cycles per run, and 500,000 coalescent simulations to approximate the expected SFS in each cycle. For each simulation replicate, the best fit was that with the smallest difference between the maximum observed and the maximum estimated likelihood.

### Demographic inference with *δaδi*

As with fastsimcoal2, demographic inference with *δaδi* (version 2.0.5) was performed on all SNP data, and separately with exonic SNPs masked, with spectra composed for each set of 10 replicates. *δaδi* was used to fit each SFS to a two-epoch demographic model. This model fits two parameters: the current population size relative to the ancestral population size (*Nu_opt_*) and the time of change relative to the ancestral population size (*τ_opt_*). For each simulation replicate, 15 starting values between −2 and 2 were drawn for *Nu* and 8 starting values between −2 and 2 were drawn for *τ*, both evenly distributed in log space. This created a total of 120 different starting parameterizations. To encourage better convergence, the maximum number of iterations for the optimizer was set to 100. The best fit for each simulation replicate was that with the lowest log-likelihood score.

### DFE inference

The gamma-distributed DFE for the simulated data was inferred using Grapes (version 1.1) with the *GammaZero* model (Galtier 2016), DFE-alpha (release 2.16; Keightley and Eyre-Walker 2007), and polyDFE (Tataru and Bataillon 2020). For Grapes and DFE-alpha, batches of 10 simulation replicates were combined to generate spectra. For analyses with Grapes, the proportion of mutations in each of the four DFE categories was estimated from the inferred gamma distribution. In addition, Grapes also infers the likelihood of a neutral model (see Supplementary File S2 for comparisons of likelihoods under the *GammaZero* and neutral models). DFE-alpha readily supplies the proportion of mutations in each of the four DFE categories via the “prop_muts_in_s_ranges” function. The mean value across each set of 10 spectra was used for comparison to the six simulated DFE models. For polyDFE, inference was performed exclusively on the equilibrium population simulations with fixed recombination rate and variable mutation rates, and performed on each simulation replicate. Within each replicate, each variable mutation rate region was provided as a separate SFS fragment in the input file, to allow polyDFE to account for mutation rate variation. As with Grapes, the proportion of mutations in each of the four DFE categories was estimated from the inferred gamma distribution.

### Calculating summary statistics

Summary statistics were calculated across 5kb sliding windows with a step size of 2,500bp, using the python implementation of libsequence (version 1.8.3) (Thornton 2003).

## Data availability

All scripts to generate and analyse simulated data, as well as to perform demographic and DFE inference, are available at the Github repository:

https://github.com/vivaksoni/rate_heterogeneity.

## Acknowledgements

We thank Parul Johri for helpful comment and discussion. This research was conducted using resources provided by Research Computing at Arizona State University (http://www.researchcomputing.asu.edu) and the Open Science Grid, which is supported by the National Science Foundation and the U.S. Department of Energy’s Office of Science. This project was funded by National Institutes of Health grant R35GM139383 to JDJ.

