## Supplementary Figure for "The effects of mutation and recombination rate heterogeneity on the inference of demography and the distribution of fitness effects"

**S1**

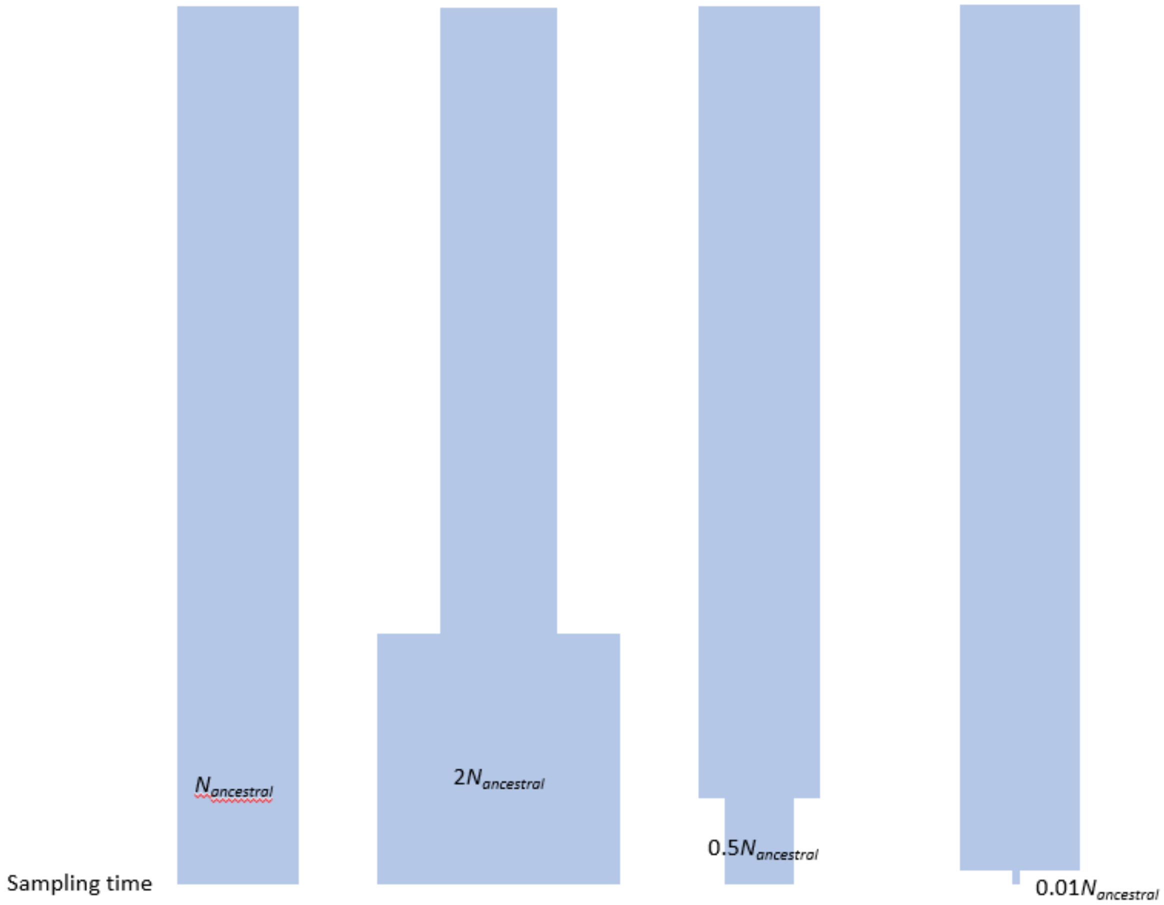


**S1:** 4 simulated demographic histories. In all scenarios a $16N_{ancestral}$ burn-in is simulated. Population size change occurs instantaneously, $N_{current}$ generations before sampling.

**S2**

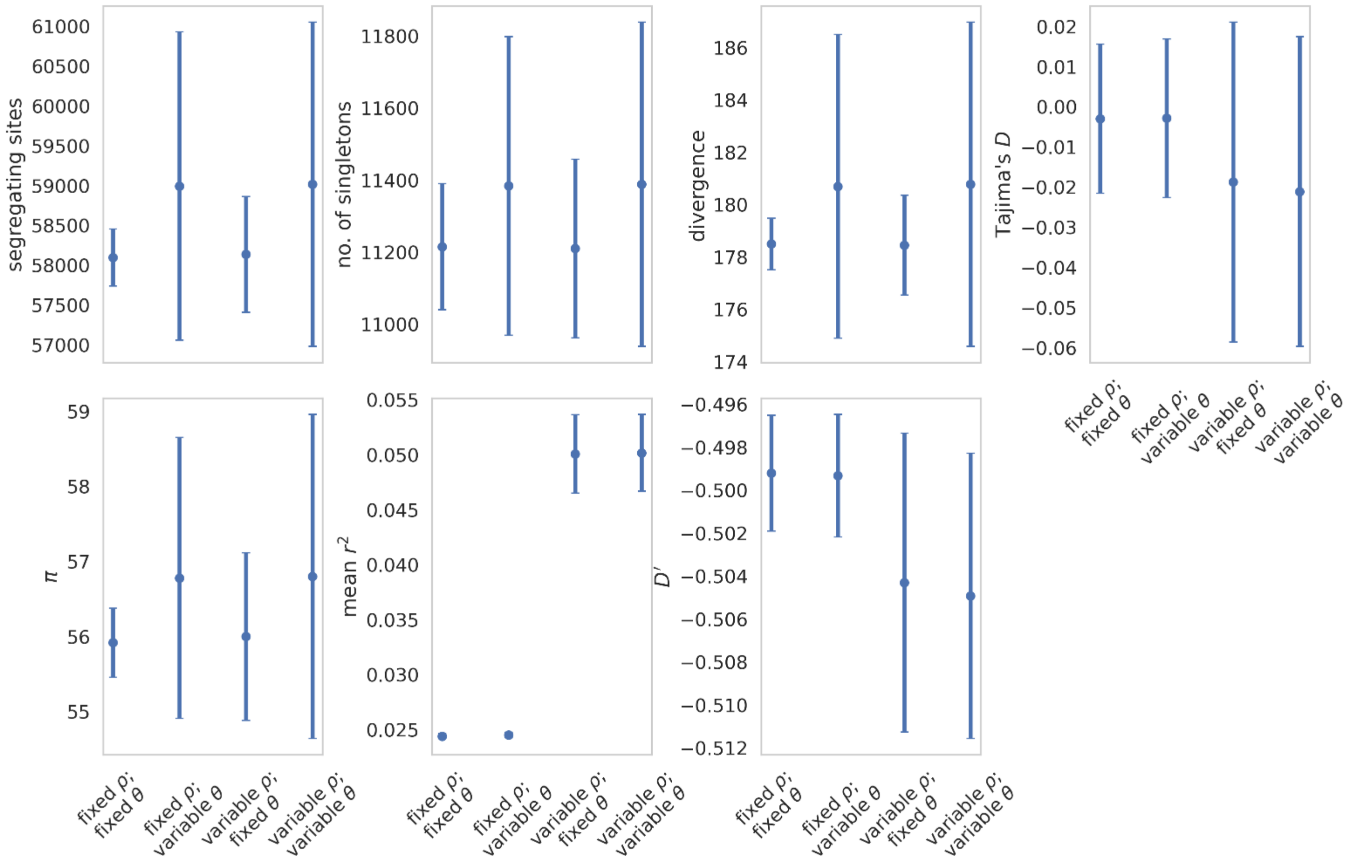


**S2:** Summary statistics for **neutrally evolving equilibrium population** under fixed and variable recombination and mutation rates. Data points represent the mean value, whilst error bars represent the standard deviation. $\theta = 2N\mu$ and $\rho = 2Nr$, where $N$ is the ancestral population size ($N_{ancestral}$), $\mu$ is the mutation rate, and $r$ is the rate of recombination. Variable rates are drawn from a uniform distribution, such that the mean rate across each simulation replicate was equal to the fixed rate, to enable fair comparisons (see the Materials and Methods section for further details).

**S3**

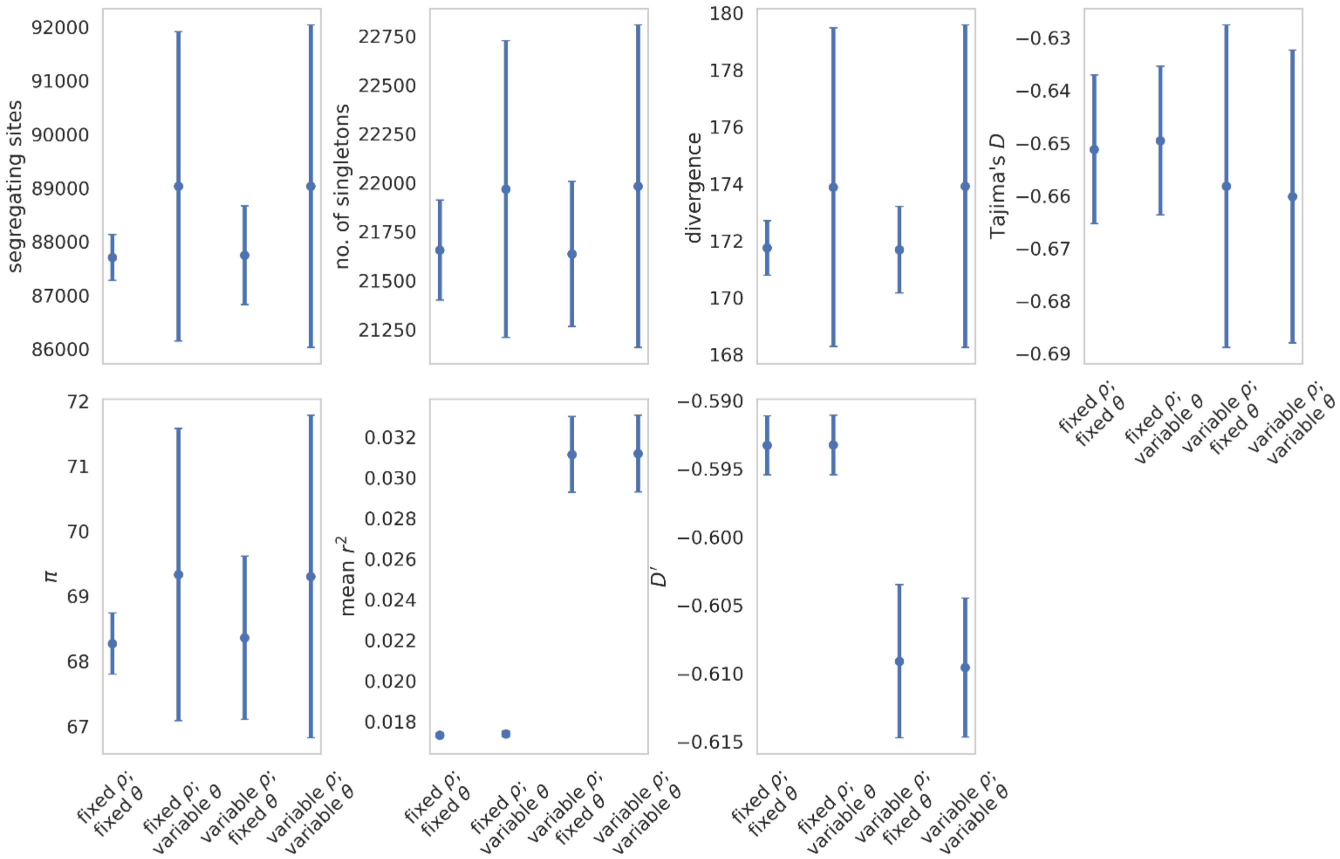


**S3:** Summary statistics for a **neutrally evolving population that has undergone population expansion** $1N_{ancestral}$ generations before sampling (**whereby $N_{current} = 2N_{ancestral}$**), under fixed and variable recombination and mutation rates. Data point represent the mean value, whilst error bars represent the standard deviation. $\theta = 2N\mu$ and $\rho = 2Nr$, where $N$ is the ancestral population size ($N_{ancestral}$), $\mu$ is the mutation rate, and $r$ is the rate of recombination. Variable rates are drawn from a uniform distribution, such that the mean rate across each simulation replicate was equal to the fixed rate, to enable fair comparisons (see the Materials and Methods section for further details).

**S4**

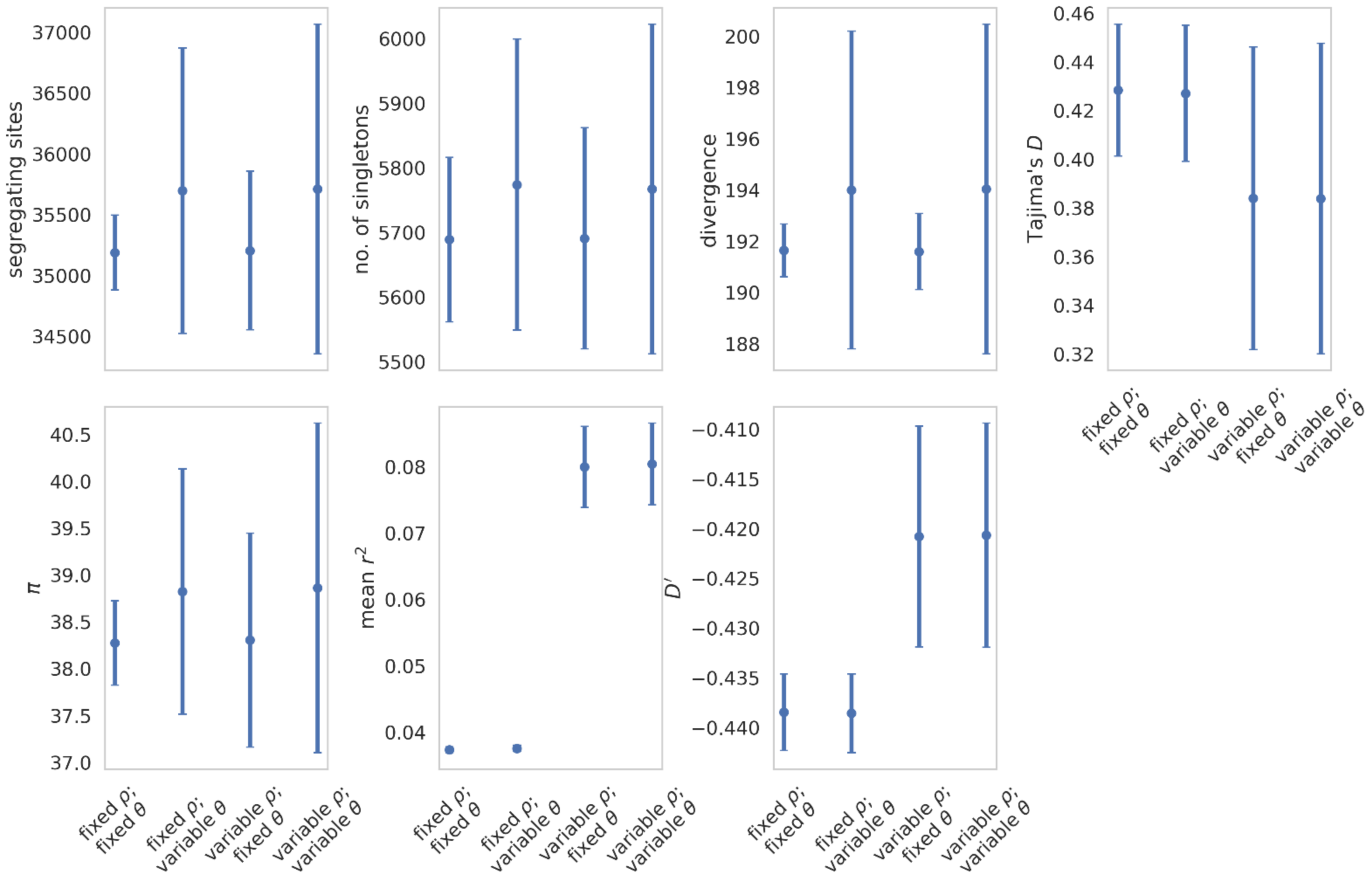


**S4:** Summary statistics for a **neutrally evolving population that has undergone population contraction** 1$N_{ancestral}$ generations before sampling (**whereby $N_{current}$ = 0.5$N_{ancestral}$**), under fixed and variable recombination and mutation rates. Data point represent the mean value, whilst error bars represent the standard deviation. $\theta = 2N\mu$ and $\rho = 2Nr$, where $N$ is the ancestral population size ($N_{ancestral}$), $\mu$ is the mutation rate, and $r$ is the rate of recombination. Variable rates are drawn from a uniform distribution, such that the mean rate across each simulation replicate was equal to the fixed rate, to enable fair comparisons (see the Materials and Methods section for further details).

**S5**

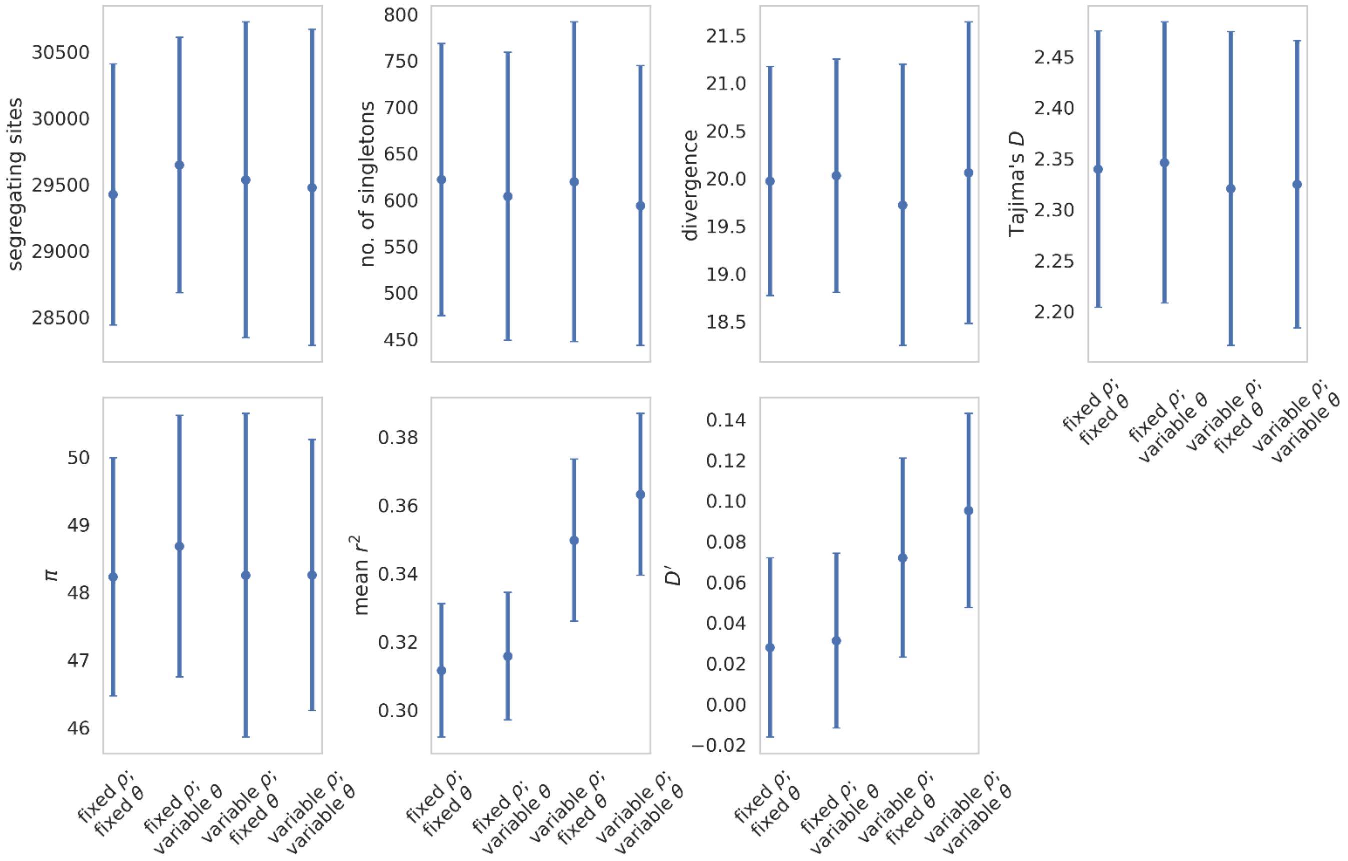


**S5:** Summary statistics for a **neutrally evolving population that has undergone population contraction** $1N_{ancestral}$ generations before sampling (**whereby** $N_{current} = 0.01N_{ancestral}$), under fixed and variable recombination and mutation rates. Data point represent the mean value, whilst error bars represent the standard deviation. $\theta = 2N\mu$ and $\rho = 2Nr$, where $N$ is the ancestral population size ($N_{ancestral}$), $\mu$ is the mutation rate, and $r$ is the rate of recombination. Variable rates are drawn from a uniform distribution, such that the mean rate across each simulation replicate was equal to the fixed rate, to enable fair comparisons (see the Materials and Methods section for further details).

**S6**

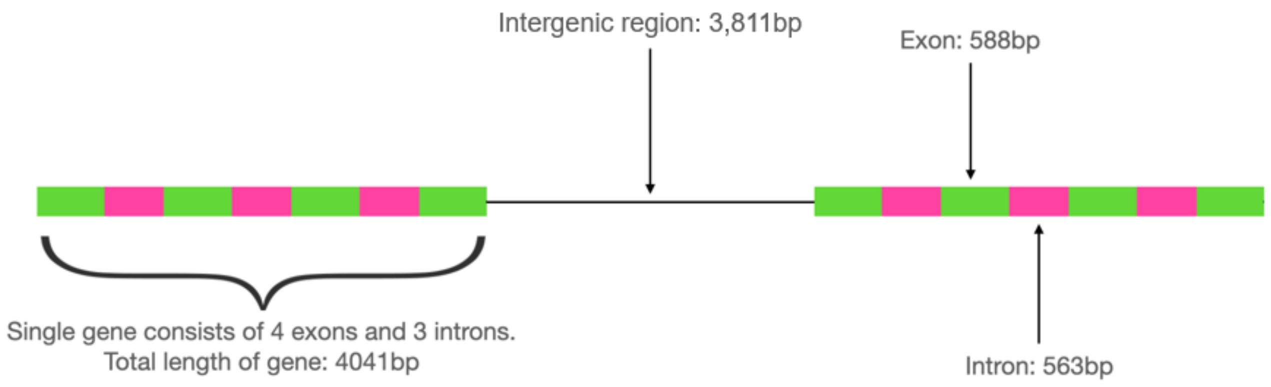


**S6:** Schematic of genic structure for DFE simulations. Each simulated region is made up of 127 genes, with each pair separated by an intergenic region.

**S7**

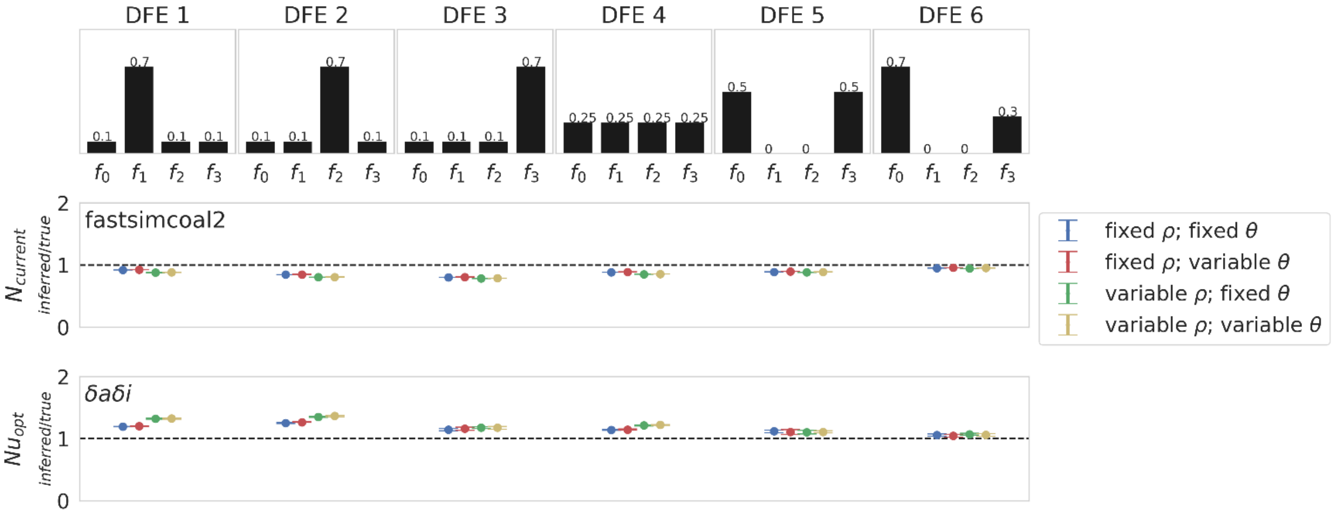


**S7:** Demographic inference results for **a single equilibrium population under multiple distributions of fitness effects (DFEs), with exonic regions unmasked**. Top row depicts the 6 discrete DFEs that were simulated. Exonic mutations were drawn from a DFE comprised of four fixed classes (Johri et al. 2020), whose frequencies were denoted by $f_i$: $f_0$ with $0 \leq 2N_{ancestral}\, s < 1$ (*i.e.*, effectively neutral mutations), $f_1$ with $1 \leq 2N_{ancestral}\, s < 10$ (*i.e.*, weakly deleterious mutations), $f_2$ with $10 \leq 2N_{ancestral}\, s < 100$ (*i.e.*, moderately deleterious mutations), and $f_3$ with $100 \leq 2N_{ancestral}\, s$ (*i.e.*, strongly deleterious mutations), where $N_{ancestral}$ was the initial population size and *s* was the reduction in fitness of the mutant homozygote relative to wild-type. Middle and bottom rows show inference results for fastsimcoal2 and *δaδI,* respectively. All inference values are scaled by the true value. As such, a value of 1 on the Y-axis indicates that the inferred parameter value matches the true parameter value (as shown by the black dashed line). For the equilibrium population model, fastsimcoal2 infers the $N_{current}$ parameter, which is the population size at time of sampling, whilst *δaδi* infers $Nu_{opt}$, which is the population size at time of sampling relative to the initial population size. Data points represent the mean inference value across 10 replicates, whilst error bars represent the standard deviation. $\theta = 2N\mu$ and $\rho = 2Nr$, where *N* is the ancestral population size ($N_{ancestral}$), $\mu$ is the per-site per-generation mutation rate, and *r* is the per-site per-generation recombination rate. Variable rates were drawn from a uniform distribution, such that the mean rate across each simulation replicate was equal to the fixed rate, to enable fair comparisons (see the Materials and Methods section for further details).

**S8**

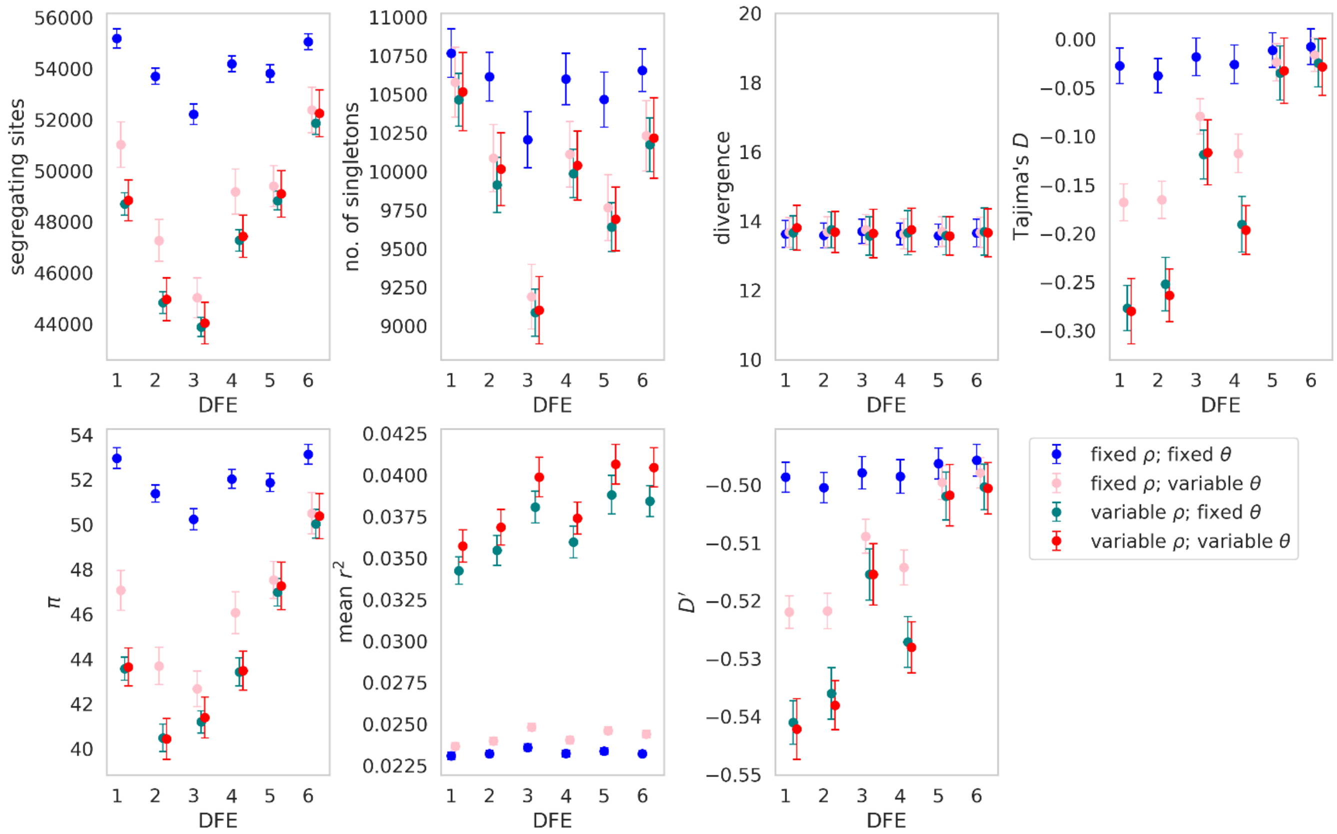


**S8:** Summary statistics for an **equilibrium population simulated under 6 different distribution of fitness effects (DFEs),** under fixed and variable recombination and mutation rates, with **exonic regions masked**. Exonic mutations were drawn from a DFE comprised of four fixed classes (Johri et al. 2020), whose frequencies were denoted by $f_i$: $f_0$, with $0 \leq 2N_{ancestral}s < 1$ (i.e., effectively neutral mutations), $f_1$, with $1 \leq 2N_{ancestral}s < 10$ (i.e., weakly deleterious mutations), $f_2$, with $10 \leq 2N_{ancestral}s < 100$ (i.e., moderately deleterious mutations), and $f_3$, with $100 \leq 2N_{ancestral}s < 2N_{ancestral}$ (i.e., strongly deleterious mutations), where $N_{ancestral}$ was the initial population size and $s$ was the reduction in fitness of the mutant homozygote relative to wild-type. These summary statistics correspond to demographic inference in Figure 2 in main text. Data point represent the mean value, whilst error bars represent the standard deviation. $\theta = 2N\mu$ and $\rho = 2Nr$, where $N$ is the ancestral population size ($N_{ancestral}$), $\mu$ is the mutation rate, and $r$ is the rate of recombination. Variable rates are drawn from a uniform distribution, such that the mean rate across each simulation replicate was equal to the fixed rate, to enable fair comparisons (see the Materials and Methods section for further details).

**S9**

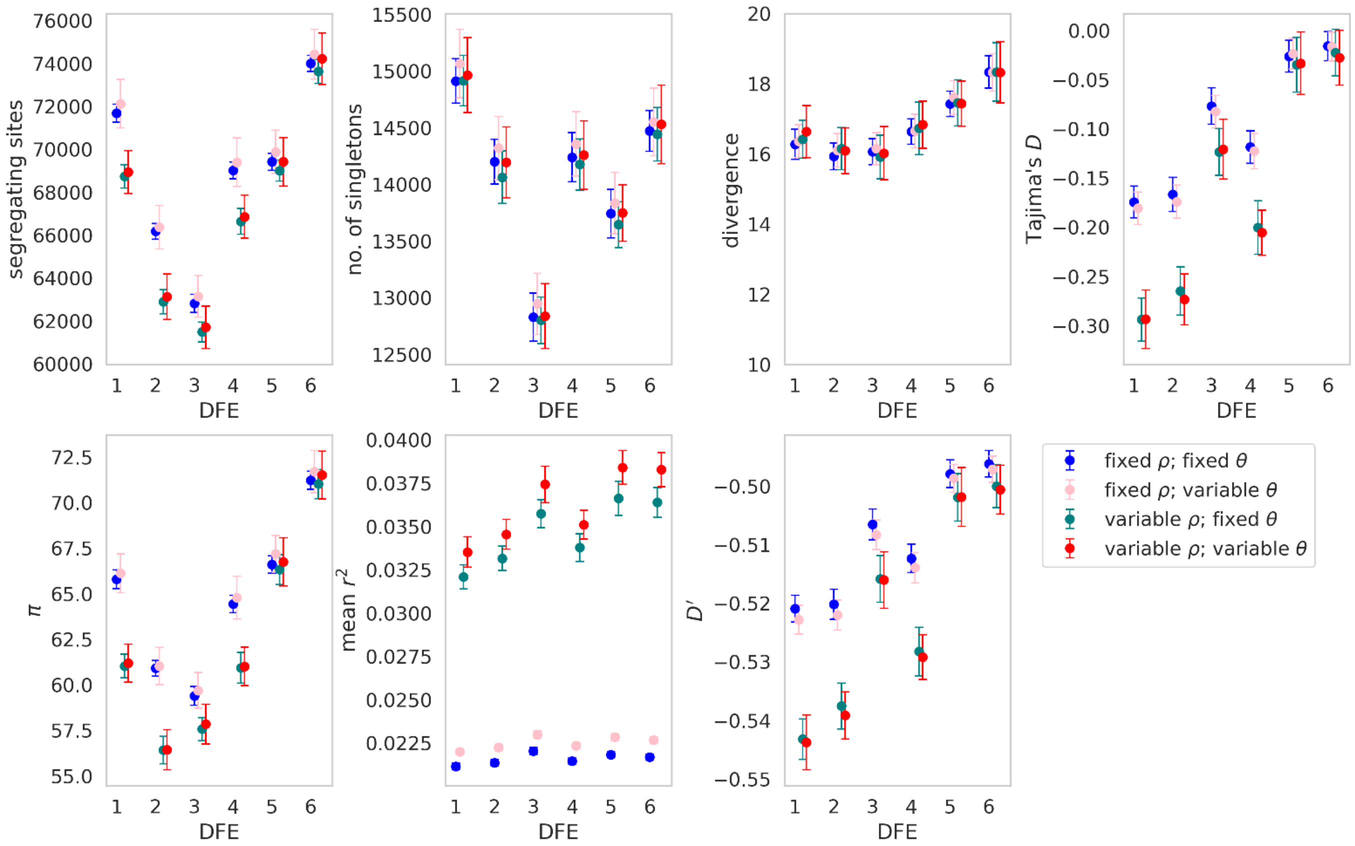


**S9:** Summary statistics for an **equilibrium population simulated under 6 different distribution of fitness effects (DFEs),** under fixed and variable recombination and mutation rates, with **no masking**. Exonic mutations were drawn from a DFE comprised of four fixed classes (Johri et al. 2020), whose frequencies were denoted by $f_i$: $f_0$, with $0 \leq 2N_{ancestral}s < 1$ (i.e., effectively neutral mutations), $f_1$, with $1 \leq 2N_{ancestral}s < 10$ (i.e., weakly deleterious mutations), $f_2$, with $10 \leq 2N_{ancestral}s < 100$ (i.e., moderately deleterious mutations), and $f_3$, with $100 \leq 2N_{ancestral}s < 2N_{ancestral}$ (i.e., strongly deleterious mutations), where $N_{ancestral}$ was the initial population size and *s* was the reduction in fitness of the mutant homozygote relative to wild-type. These summary statistics correspond to demographic inference in Supplementary Figure S7 in main text. Data point represent the mean value, whilst error bars represent the standard deviation. $\theta = 2N\mu$ and $\rho = 2Nr$, where *N* is the ancestral population size ($N_{ancestral}$), $\mu$ is the mutation rate, and *r* is the rate of recombination. Variable rates are drawn from a uniform distribution, such that the mean rate across each simulation replicate was equal to the fixed rate, to enable fair comparisons (see the Materials and Methods section for further details).

**S10**

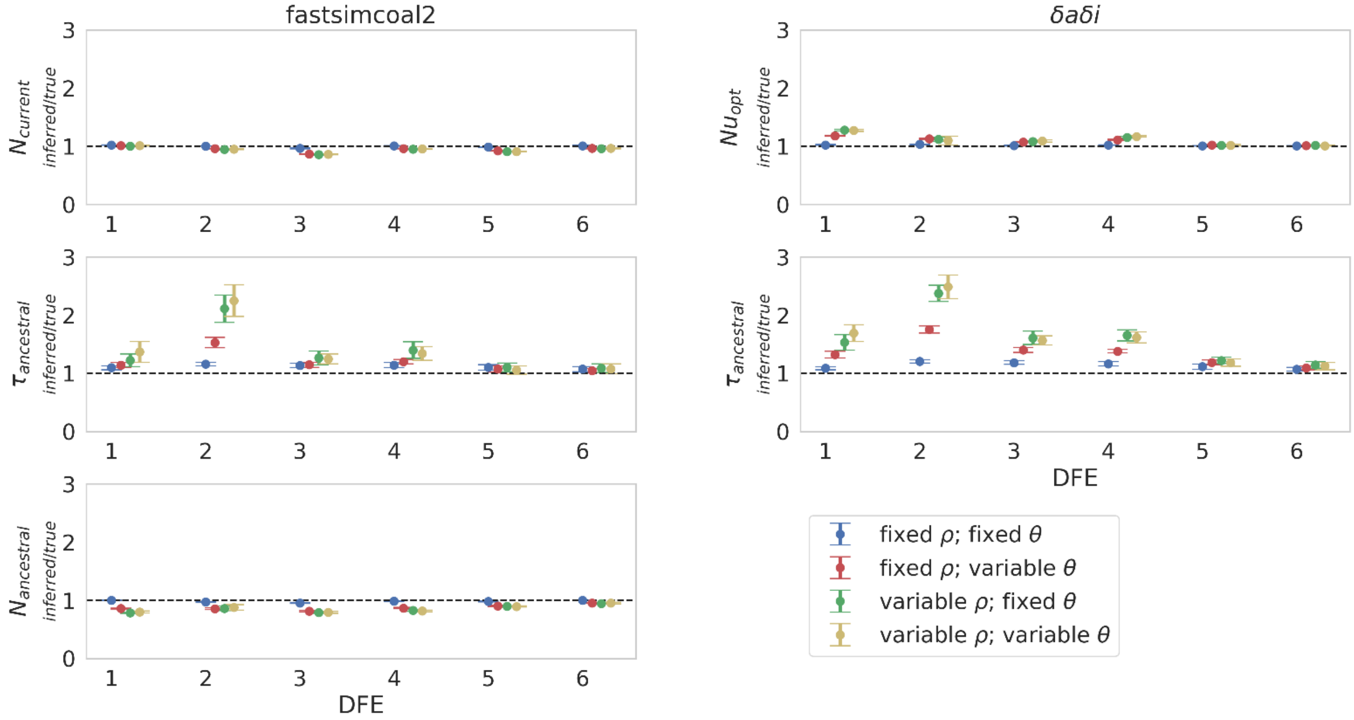


**S10:** Demographic inference results for a **population contraction (such that $N_{current}$ = 0.5$N_{ancestral}$)** under fixed and variable recombination and mutation rates, across the 6 simulated distributions of fitness effects (DFEs), with **exonic regions masked**. Population size change occurred 1$N_{current}$ generations before sampling, where $N_{current}$ is the population size at time of sampling. Exonic mutations were drawn from a DFE comprised of four fixed classes (Johri et al. 2020), whose frequencies were denoted by $f_i$: $f_0$ with $0 \leq 2N_{ancestral}\, s < 1$ (*i.e.*, effectively neutral mutations), $f_1$ with $1 \leq 2N_{ancestral}\, s < 10$ (*i.e.*, weakly deleterious mutations), $f_2$ with $10 \leq 2N_{ancestral}\, s < 100$ (*i.e.*, moderately deleterious mutations), and $f_3$ with $100 \leq 2N_{ancestral}\, s$ (*i.e.*, strongly deleterious mutations), where $N_{ancestral}$ was the initial population size and *s* was the reduction in fitness of the mutant homozygote relative to wild-type. fastsimcoal2 infers three parameters: $N_{current}$; $N_{ancestral}$; and $\tau_{ancestral}$ (the time of population size change in $N_{current}$ generations). *δaδI* infers two parameters: $Nu_{opt}$ (the population size at time of sampling relative to the initial population size); and $\tau_{opt}$ (the time of population size change relative to the initial population size). Data points represent the mean inference value across 10 replicates, whilst error bars represent the standard deviation. $\theta = 2N\mu$ and $\rho = 2Nr$, where *N* is the ancestral population size ($N_{ancestral}$), $\mu$ is the per-site per-generation mutation rate, and *r* is the per-site per-generation recombination rate. Variable rates are drawn from a uniform distribution, such that the mean rate across each simulation replicate was equal to the fixed rate, to enable fair comparisons (see the Materials and Methods section for further details).

**S11**

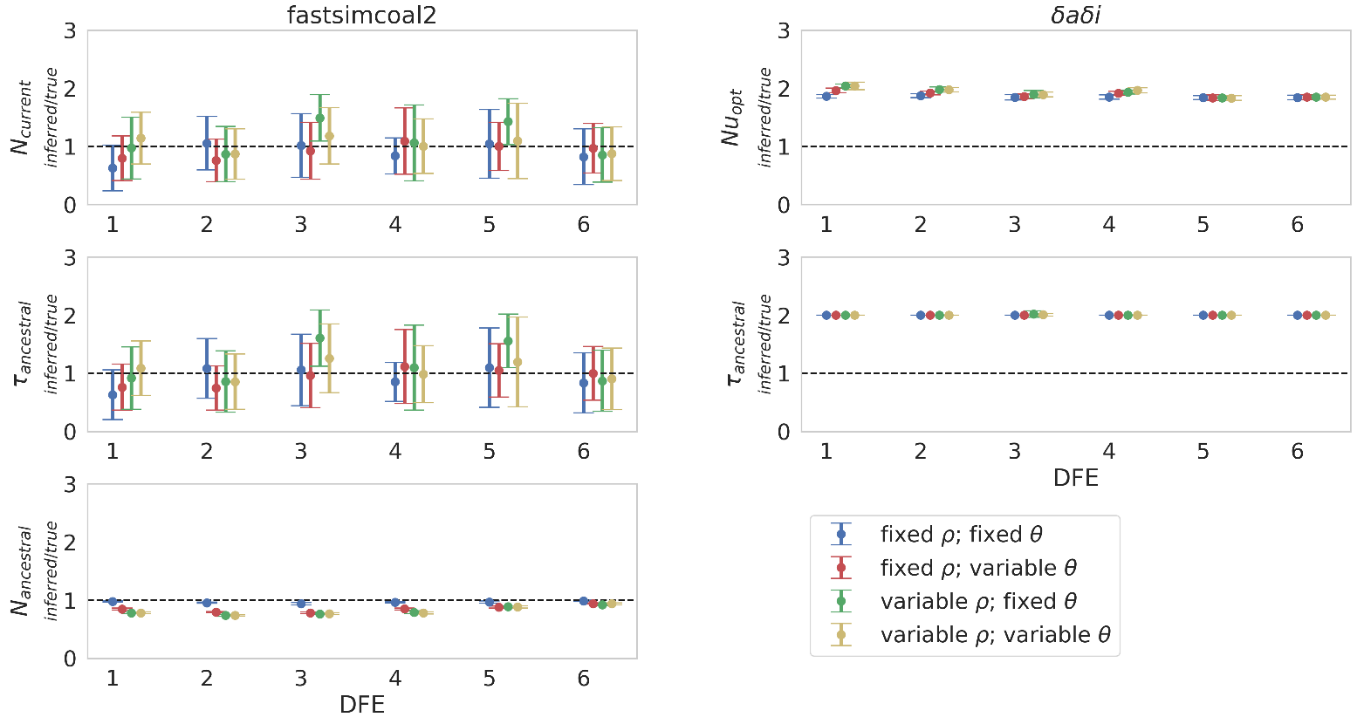


**S11:** Demographic inference results for a **population contraction (such that $N_{current}$ = 0.01$N_{ancestral}$)** under fixed and variable recombination and mutation rates, across the 6 simulated distributions of fitness effects (DFEs), with **exonic regions masked**. Population size change occurred 1$N_{current}$ generations before sampling, where $N_{current}$ is the population size at time of sampling. Exonic mutations were drawn from a DFE comprised of four fixed classes (Johri et al. 2020), whose frequencies were denoted by $f_i$: $f_0$ with $0 \leq 2N_{ancestral}\, s < 1$ (*i.e.*, effectively neutral mutations), $f_1$ with $1 \leq 2N_{ancestral}\, s < 10$ (*i.e.*, weakly deleterious mutations), $f_2$ with $10 \leq 2N_{ancestral}\, s < 100$ (*i.e.*, moderately deleterious mutations), and $f_3$ with $100 \leq 2N_{ancestral}\, s$ (*i.e.*, strongly deleterious mutations), where $N_{ancestral}$ was the initial population size and $s$ was the reduction in fitness of the mutant homozygote relative to wild-type. fastsimcoal2 infers three parameters: $N_{current}$; $N_{ancestral}$; and $\tau_{ancestral}$ (the time of population size change in $N_{current}$ generations). *δaδi* infers two parameters: $Nu_{opt}$ (the population size at time of sampling relative to the initial population size); and $\tau_{opt}$ (the time of population size change relative to the initial population size). Data points represent the mean inference value across 10 replicates, whilst error bars represent the standard deviation. $\theta = 2N\mu$ and $\rho = 2Nr$, where $N$ is the ancestral population size ($N_{ancestral}$), $\mu$ is the per-site per-generation mutation rate, and $r$ is the per-site per-generation recombination rate. Variable rates are drawn from a uniform distribution, such that the mean rate across each simulation replicate was equal to the fixed rate, to enable fair comparisons (see the Materials and Methods section for further details).

**S12**

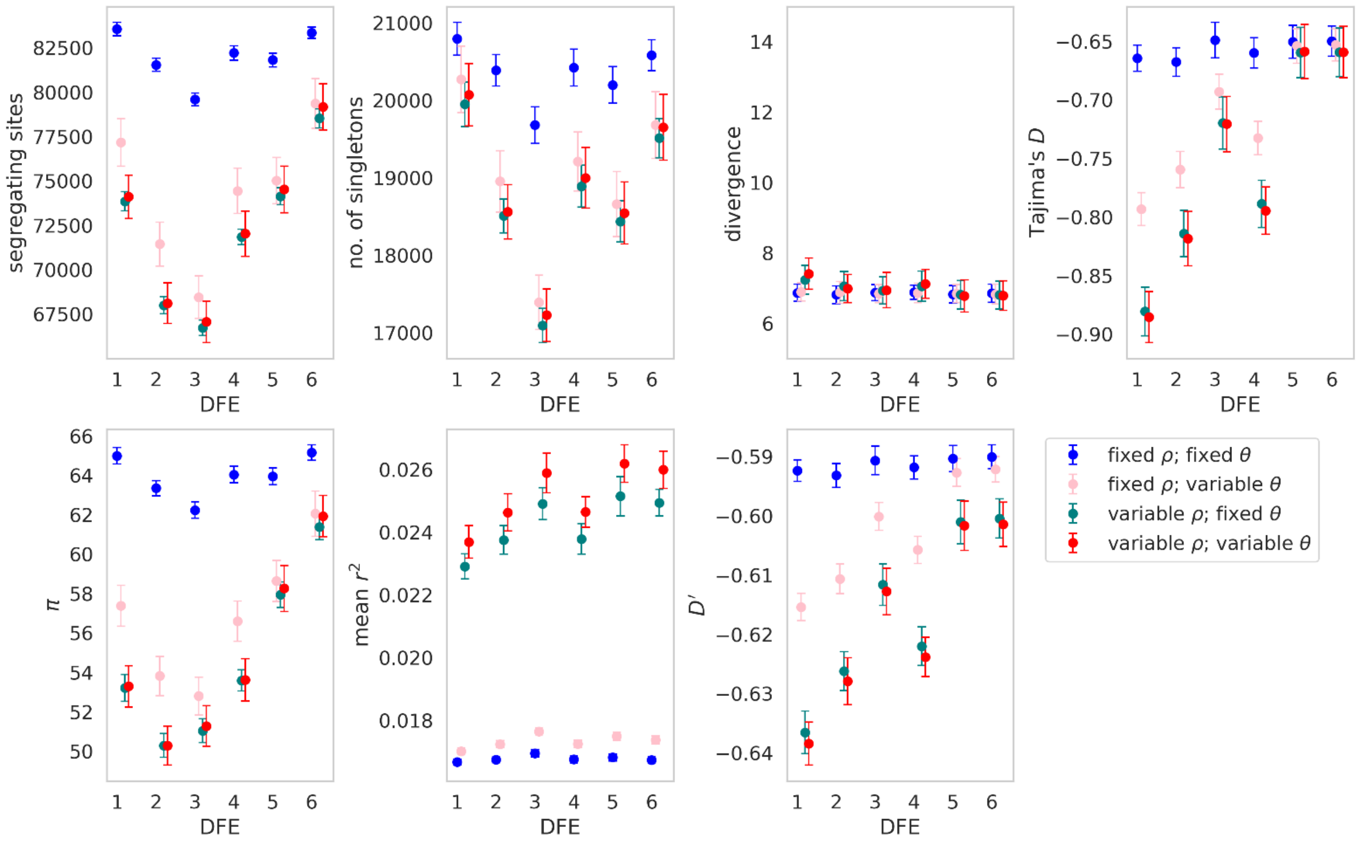


**S12:** Summary statistics for a **population expansion (such that $N_{current}$ = 2$N_{ancestral}$) simulated under 6 different distribution of fitness effects (DFEs),** under fixed and variable recombination and mutation rates, with **exonic regions masked**. Exonic mutations were drawn from a DFE comprised of four fixed classes (Johri et al. 2020), whose frequencies were denoted by $f_i$: $f_0$, with $0 \leq 2N_{ancestral}s < 1$ (i.e., effectively neutral mutations), $f_1$, with $1 \leq 2N_{ancestral}s < 10$ (i.e., weakly deleterious mutations), $f_2$, with $10 \leq 2N_{ancestral}s < 100$ (i.e., moderately deleterious mutations), and $f_3$, with $100 \leq 2N_{ancestral}s < 2N_{ancestral}$ (i.e., strongly deleterious mutations), where $N_{ancestral}$ was the initial population size and *s* was the reduction in fitness of the mutant homozygote relative to wild-type. These summary statistics correspond to demographic inference in Figure 3 in main text. Data point represent the mean value, whilst error bars represent the standard deviation. $\theta = 2N\mu$ and $\rho = 2Nr$, where *N* is the ancestral population size ($N_{ancestral}$), $\mu$ is the mutation rate, and *r* is the rate of recombination. Variable rates are drawn from a uniform distribution, such that the mean rate across each simulation replicate was equal to the fixed rate, to enable fair comparisons (see the Materials and Methods section for further details).

**S13**

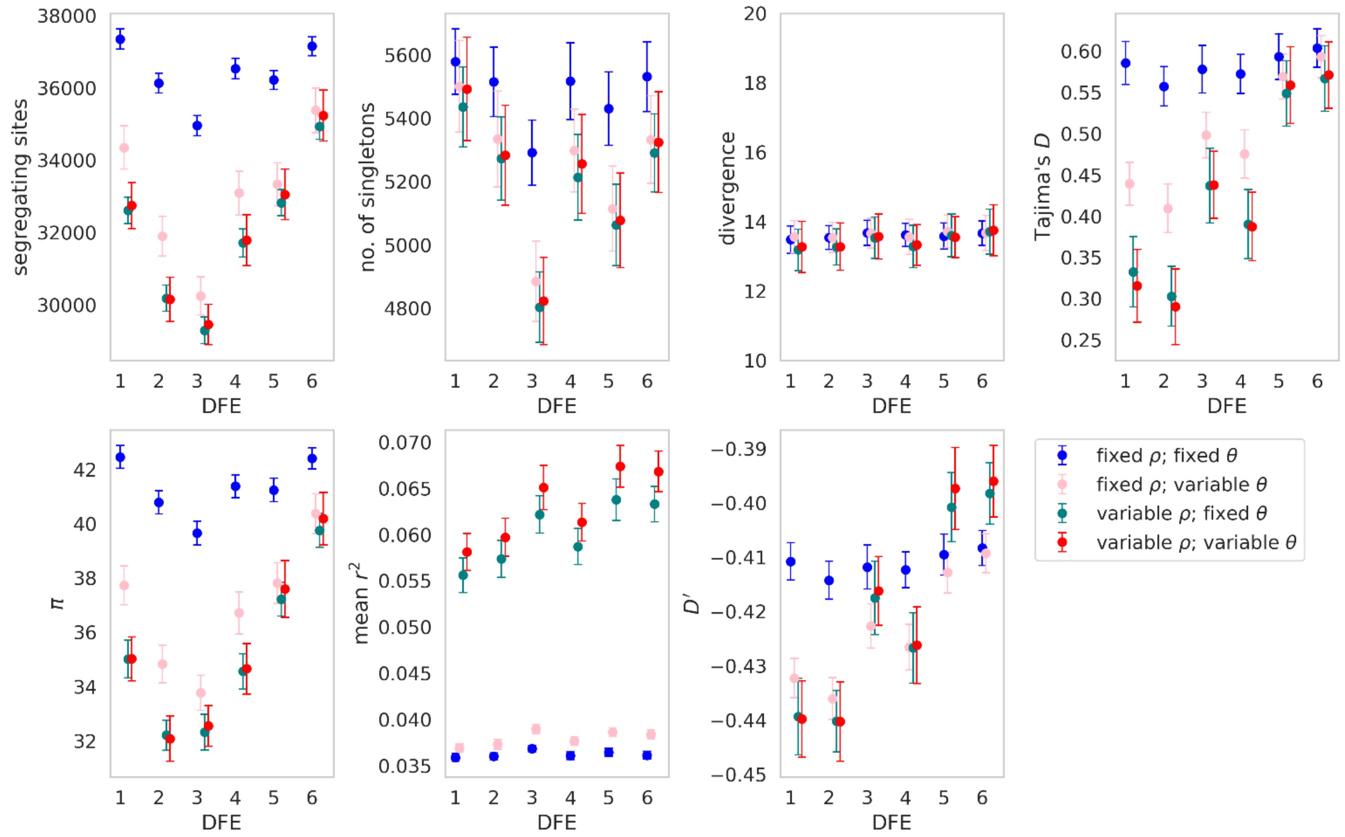


**S13:** Summary statistics for a **population contraction (such that $N_{current}$ = 0.5$N_{ancestral}$) simulated under 6 different distribution of fitness effects (DFEs),** under fixed and variable recombination and mutation rates, with **exonic regions masked**. Exonic mutations were drawn from a DFE comprised of four fixed classes (Johri et al. 2020), whose frequencies were denoted by $f_i$: $f_0$, with $0 \leq 2N_{ancestral}s < 1$ (i.e., effectively neutral mutations), $f_1$, with $1 \leq 2N_{ancestral}s < 10$ (i.e., weakly deleterious mutations), $f_2$, with $10 \leq 2N_{ancestral}s < 100$ (i.e., moderately deleterious mutations), and $f_3$, with $100 \leq 2N_{ancestral}s < 2N_{ancestral}$ (i.e., strongly deleterious mutations), where $N_{ancestral}$ was the initial population size and $s$ was the reduction in fitness of the mutant homozygote relative to wild-type. Data point represent the mean value, whilst error bars represent the standard deviation. $\theta = 2N\mu$ and $\rho = 2Nr$, where $N$ is the ancestral population size ($N_{ancestral}$), $\mu$ is the mutation rate, and $r$ is the rate of recombination. Variable rates are drawn from a uniform distribution, such that the mean rate across each simulation replicate was equal to the fixed rate, to enable fair comparisons (see the Materials and Methods section for further details).

## S14

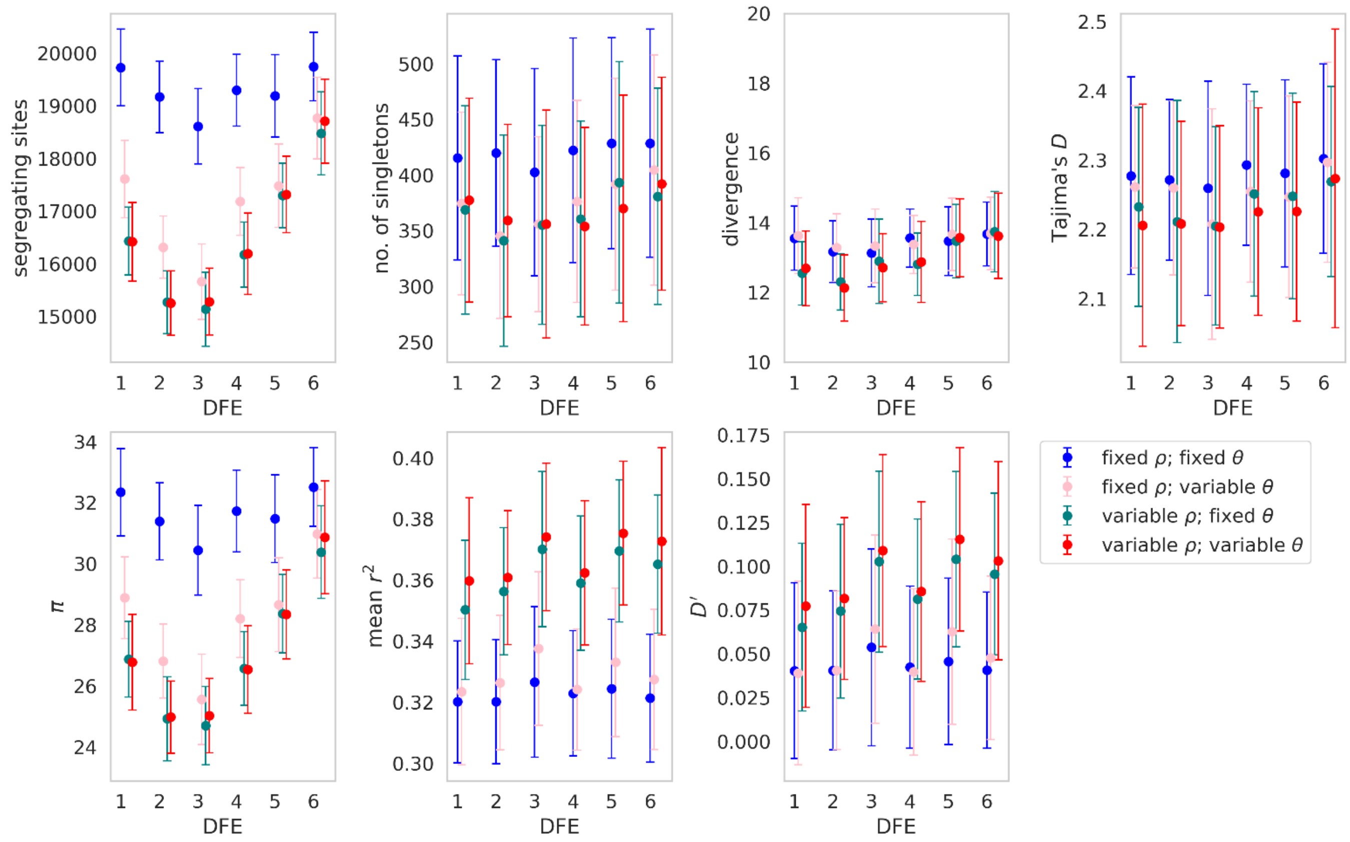


**S14:** Summary statistics for a **population contraction (such that $N_{current}$ = 0.01$N_{ancestral}$) simulated under 6 different distribution of fitness effects (DFEs),** under fixed and variable recombination and mutation rates, with **exonic regions masked**. Exonic mutations were drawn from a DFE comprised of four fixed classes (Johri et al. 2020), whose frequencies were denoted by $f_i$: $f_0$, with $0 \leq 2N_{ancestral}s < 1$ (i.e., effectively neutral mutations), $f_1$, with $1 \leq 2N_{ancestral}s < 10$ (i.e., weakly deleterious mutations), $f_2$, with $10 \leq 2N_{ancestral}s < 100$ (i.e., moderately deleterious mutations), and $f_3$, with $100 \leq 2N_{ancestral}s < 2N_{ancestral}$ (i.e., strongly deleterious mutations), where $N_{ancestral}$ was the initial population size and $s$ was the reduction in fitness of the mutant homozygote relative to wild-type. Data point represent the mean value, whilst error bars represent the standard deviation. $\theta = 2N\mu$ and $\rho = 2Nr$, where $N$ is the ancestral population size ($N_{ancestral}$), $\mu$ is the mutation rate, and $r$ is the rate of recombination. Variable rates are drawn from a uniform distribution, such that the mean rate across each simulation replicate was equal to the fixed rate, to enable fair comparisons (see the Materials and Methods section for further details).

**S15**

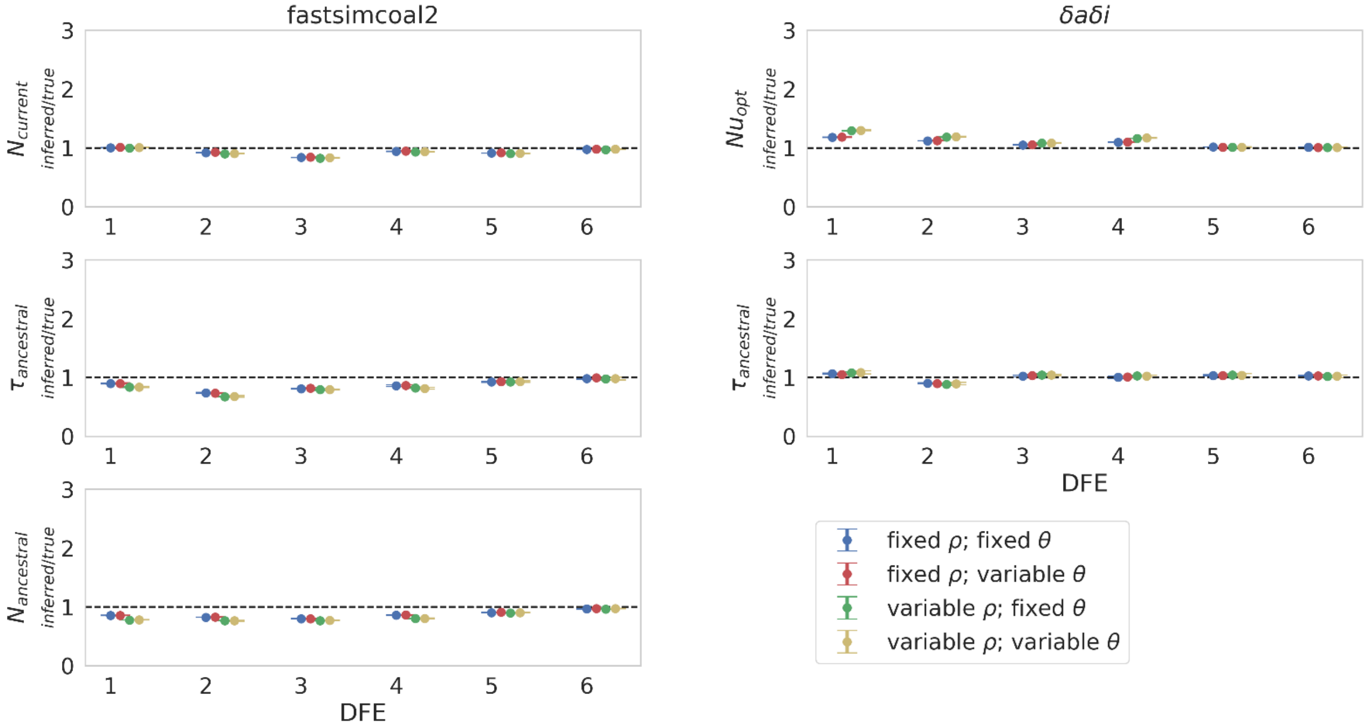


**S15:** Demographic inference results for a **population expansion (such that $N_{current}$ = 2$N_{ancestral}$)** under fixed and variable recombination and mutation rates, across the 6 simulated distributions of fitness effects (DFEs), with **exonic regions unmasked**. Population size change occurred 1$N_{current}$ generations before sampling, where $N_{current}$ is the population size at time of sampling. Exonic mutations were drawn from a DFE comprised of four fixed classes (Johri et al. 2020), whose frequencies were denoted by $f_i$: $f_0$ with $0 \leq 2N_{ancestral}\, s < 1$ (*i.e.*, effectively neutral mutations), $f_1$ with $1 \leq 2N_{ancestral}\, s < 10$ (*i.e.*, weakly deleterious mutations), $f_2$ with $10 \leq 2N_{ancestral}\, s < 100$ (*i.e.*, moderately deleterious mutations), and $f_3$ with $100 \leq 2N_{ancestral}\, s$ (*i.e.*, strongly deleterious mutations), where $N_{ancestral}$ was the initial population size and *s* was the reduction in fitness of the mutant homozygote relative to wild-type. Fastsimcoal2 infers three parameters: $N_{current}$; $N_{ancestral}$; and $\tau_{ancestral}$ (the time of population size change in $N_{current}$ generations). *δaδi* infers two parameters: $Nu_{opt}$ (the population size at time of sampling relative to the initial population size); and $\tau_{opt}$ (the time of population size change relative to the initial population size). Data points represent the mean inference value across 10 replicates, whilst error bars represent the standard deviation. $\theta = 2N\mu$ and $\rho = 2Nr$, where *N* is the ancestral population size ($N_{ancestral}$), $\mu$ is the per-site per-generation mutation rate, and *r* is the per-site per-generation recombination rate. Variable rates are drawn from a uniform distribution, such that the mean rate across each simulation replicate was equal to the fixed rate, to enable fair comparisons (see the Materials and Methods section for further details).

**S16**

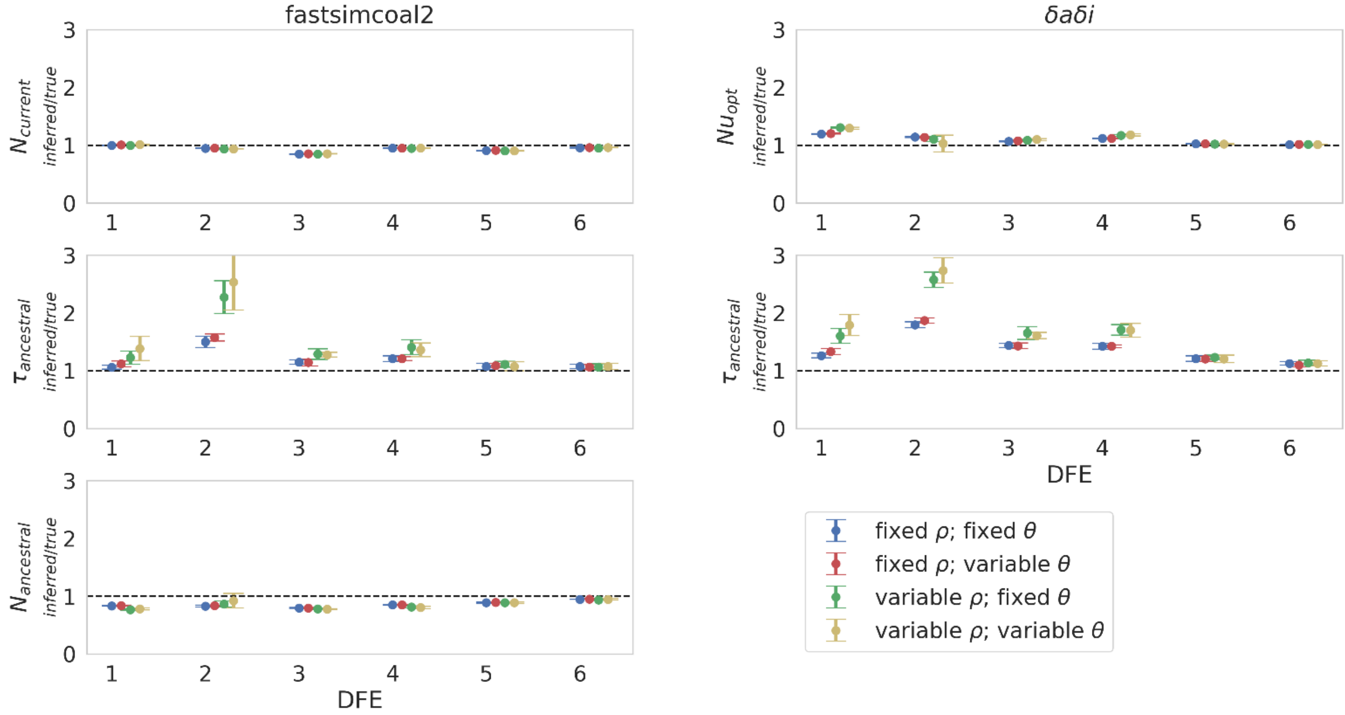


**S16:** Demographic inference results for a **population contraction (such that $N_{current}$ = 0.5$N_{ancestral}$)** under fixed and variable recombination and mutation rates, across the 6 simulated distributions of fitness effects (DFEs), with **exonic regions unmasked**. Population size change occurred 1$N_{current}$ generations before sampling, where $N_{current}$ is the population size at time of sampling. Exonic mutations were drawn from a DFE comprised of four fixed classes (Johri et al. 2020), whose frequencies were denoted by $f_i$: $f_0$ with $0 \leq 2N_{ancestral}\, s < 1$ (*i.e.*, effectively neutral mutations), $f_1$ with $1 \leq 2N_{ancestral}\, s < 10$ (*i.e.*, weakly deleterious mutations), $f_2$ with $10 \leq 2N_{ancestral}\, s < 100$ (*i.e.*, moderately deleterious mutations), and $f_3$ with $100 \leq 2N_{ancestral}\, s$ (*i.e.*, strongly deleterious mutations), where $N_{ancestral}$ was the initial population size and *s* was the reduction in fitness of the mutant homozygote relative to wild-type. Fastsimcoal2 infers three parameters: $N_{current}$; $N_{ancestral}$; and $\tau_{ancestral}$ (the time of population size change in $N_{current}$ generations). *δaδI* infers two parameters: $Nu_{opt}$ (the population size at time of sampling relative to the initial population size); and $\tau_{opt}$ (the time of population size change relative to the initial population size). Data points represent the mean inference value across 10 replicates, whilst error bars represent the standard deviation. $\theta = 2N\mu$ and $\rho = 2Nr$, where *N* is the ancestral population size ($N_{ancestral}$), $\mu$ is the per-site per-generation mutation rate, and *r* is the per-site per-generation recombination rate. Variable rates are drawn from a uniform distribution, such that the mean rate across each simulation replicate was equal to the fixed rate, to enable fair comparisons (see the Materials and Methods section for further details).

**S17**

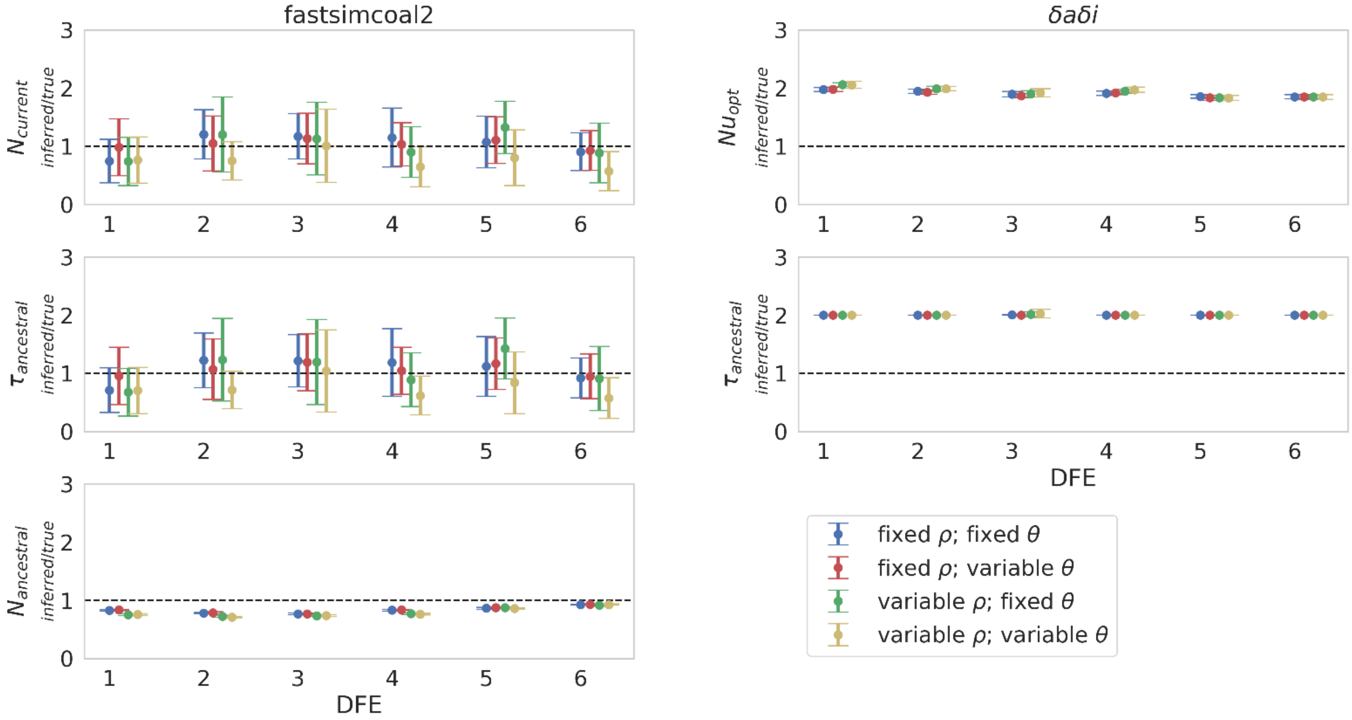


**S17:** Demographic inference results for a **population contraction (such that $N_{current}$ = 0.01$N_{ancestral}$)** under fixed and variable recombination and mutation rates, across the 6 simulated distributions of fitness effects (DFEs), with **exonic regions unmasked**. Population size change occurred 1$N_{current}$ generations before sampling, where $N_{current}$ is the population size at time of sampling. Exonic mutations were drawn from a DFE comprised of four fixed classes (Johri et al. 2020), whose frequencies were denoted by $f_i$: $f_0$ with $0 \leq 2N_{ancestral}\, s < 1$ (*i.e.*, effectively neutral mutations), $f_1$ with $1 \leq 2N_{ancestral}\, s < 10$ (*i.e.*, weakly deleterious mutations), $f_2$ with $10 \leq 2N_{ancestral}\, s < 100$ (*i.e.*, moderately deleterious mutations), and $f_3$ with $100 \leq 2N_{ancestral}\, s$ (*i.e.*, strongly deleterious mutations), where $N_{ancestral}$ was the initial population size and *s* was the reduction in fitness of the mutant homozygote relative to wild-type. Fastsimcoal2 infers three parameters: $N_{current}$; $N_{ancestral}$; and $\tau_{ancestral}$ (the time of population size change in $N_{current}$ generations). *δaδi* infers two parameters: $Nu_{opt}$ (the population size at time of sampling relative to the initial population size); and $\tau_{opt}$ (the time of population size change relative to the initial population size). Data points represent the mean inference value across 10 replicates, whilst error bars represent the standard deviation. $\theta = 2N\mu$ and $\rho = 2Nr$, where *N* is the ancestral population size ($N_{ancestral}$), $\mu$ is the per-site per-generation mutation rate, and *r* is the per-site per-generation recombination rate. Variable rates are drawn from a uniform distribution, such that the mean rate across each simulation replicate was equal to the fixed rate, to enable fair comparisons (see the Materials and Methods section for further details).

**S18**

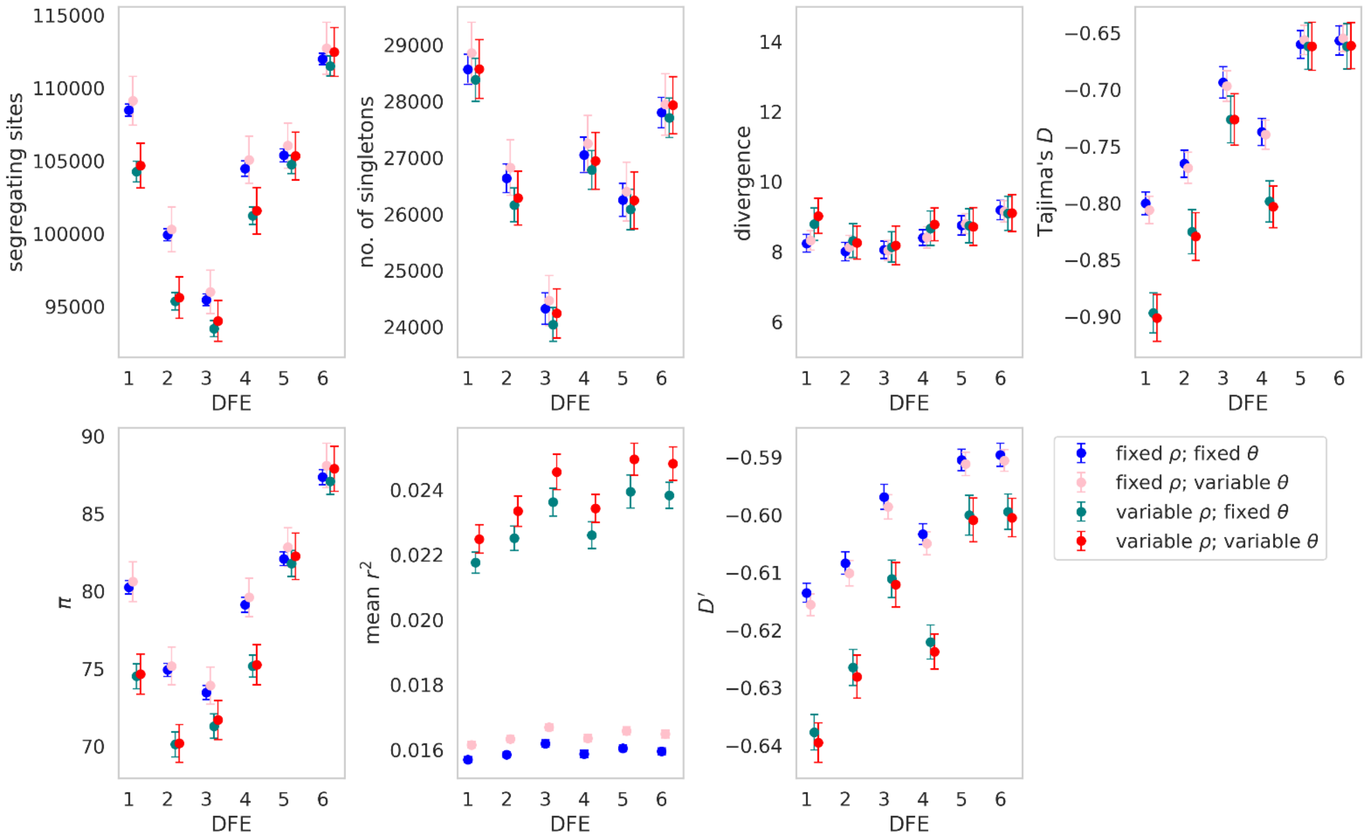


**S18:** Summary statistics for a **population expansion (such that $N_{current}$ = 2$N_{ancestral}$) simulated under 6 different distribution of fitness effects (DFEs),** under fixed and variable recombination and mutation rates, with **exonic regions unmasked**. Exonic mutations were drawn from a DFE comprised of four fixed classes (Johri et al. 2020), whose frequencies were denoted by $f_i$: $f_0$, with $0 \leq 2N_{ancestral}s < 1$ (i.e., effectively neutral mutations), $f_1$, with $1 \leq 2N_{ancestral}s < 10$ (i.e., weakly deleterious mutations), $f_2$, with $10 \leq 2N_{ancestral}s < 100$ (i.e., moderately deleterious mutations), and $f_3$, with $100 \leq 2N_{ancestral}s < 2N_{ancestral}$ (i.e., strongly deleterious mutations), where $N_{ancestral}$ was the initial population size and $s$ was the reduction in fitness of the mutant homozygote relative to wild-type. Data point represent the mean value, whilst error bars represent the standard deviation. $\theta = 2N\mu$ and $\rho = 2Nr$, where $N$ is the ancestral population size ($N_{ancestral}$), $\mu$ is the mutation rate, and $r$ is the rate of recombination. Variable rates are drawn from a uniform distribution, such that the mean rate across each simulation replicate was equal to the fixed rate, to enable fair comparisons (see the Materials and Methods section for further details).

## S19

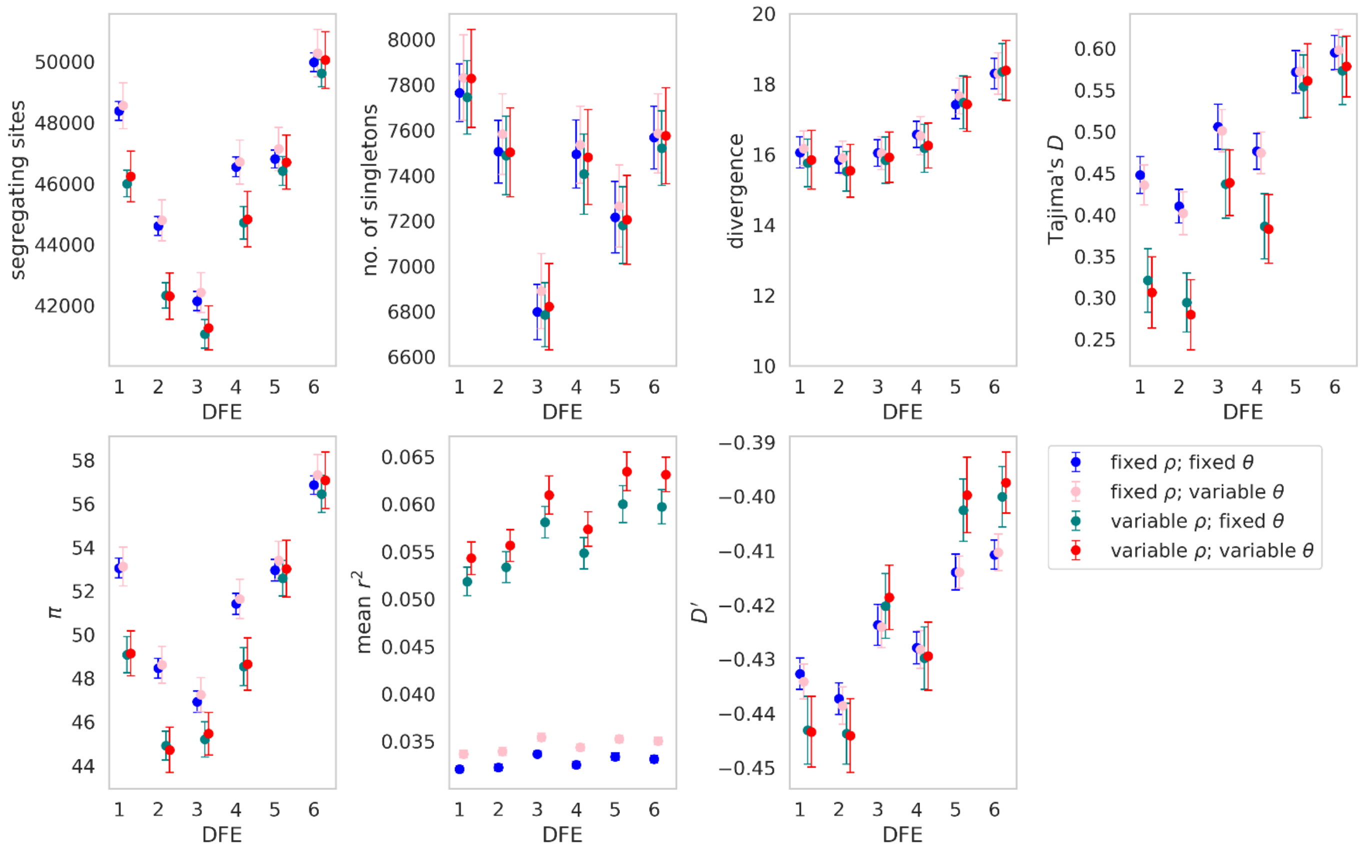


**S19:** Summary statistics for a **population contraction (such that $N_{current}$ = 0.1$N_{ancestral}$) simulated under 6 different distribution of fitness effects (DFEs),** under fixed and variable recombination and mutation rates, with **exonic regions unmasked**. Exonic mutations were drawn from a DFE comprised of four fixed classes (Johri et al. 2020), whose frequencies were denoted by $f_i$: $f_0$, with $0 \leq 2N_{ancestral}s < 1$ (i.e., effectively neutral mutations), $f_1$, with $1 \leq 2N_{ancestral}s < 10$ (i.e., weakly deleterious mutations), $f_2$, with $10 \leq 2N_{ancestral}s < 100$ (i.e., moderately deleterious mutations), and $f_3$, with $100 \leq 2N_{ancestral}s < 2N_{ancestral}$ (i.e., strongly deleterious mutations), where $N_{ancestral}$ was the initial population size and $s$ was the reduction in fitness of the mutant homozygote relative to wild-type. Data point represent the mean value, whilst error bars represent the standard deviation. $\theta = 2N\mu$ and $\rho = 2Nr$, where $N$ is the ancestral population size ($N_{ancestral}$), $\mu$ is the mutation rate, and $r$ is the rate of recombination. Variable rates are drawn from a uniform distribution, such that the mean rate across each simulation replicate was equal to the fixed rate, to enable fair comparisons (see the Materials and Methods section for further details).

**S20**

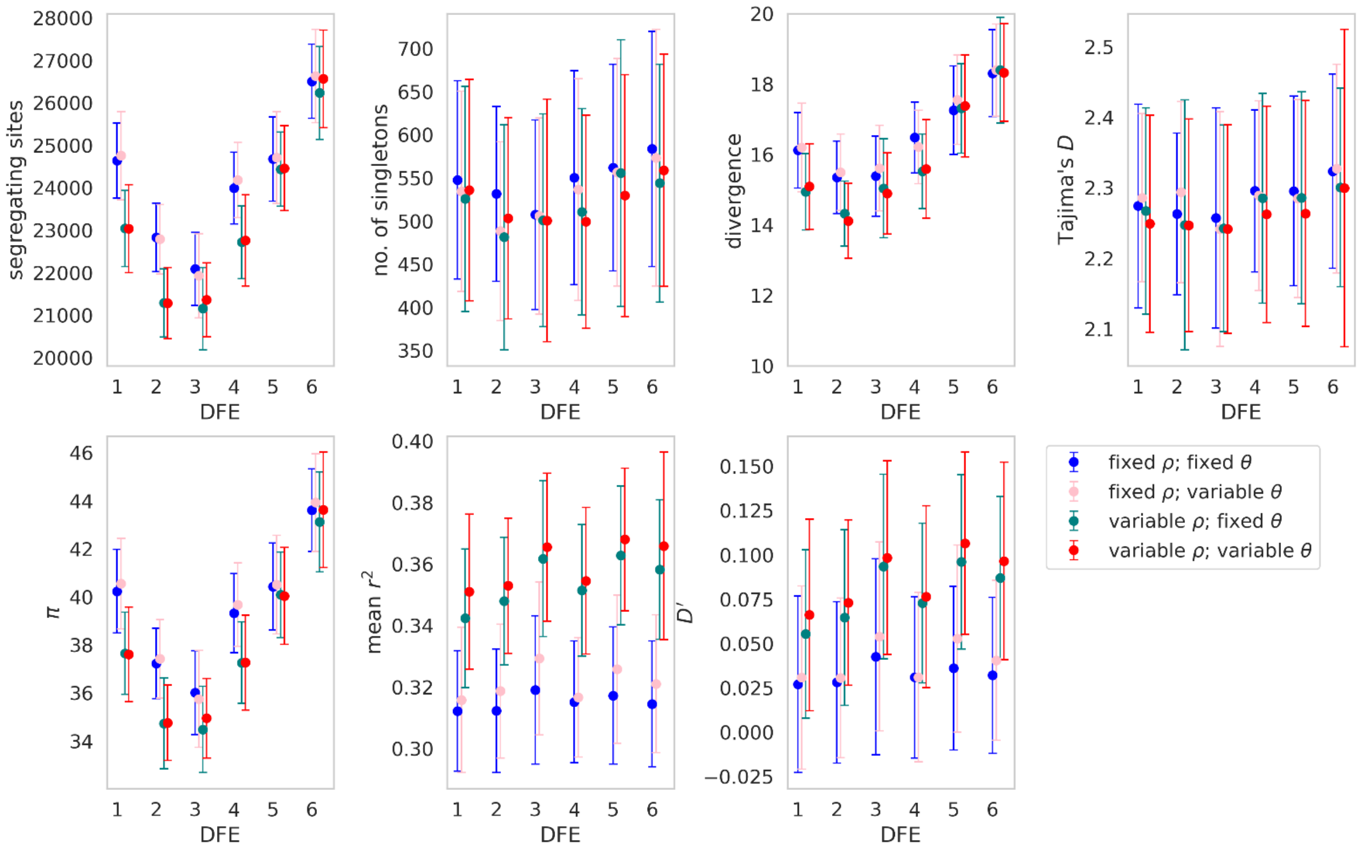


**S20:** Summary statistics for a **population contraction (such that $N_{current}$ = 0.01$N_{ancestral}$) simulated under 6 different distribution of fitness effects (DFEs),** under fixed and variable recombination and mutation rates, with **exonic regions unmasked**. Exonic mutations were drawn from a DFE comprised of four fixed classes (Johri et al. 2020), whose frequencies were denoted by $f_i$: $f_0$, with $0 \leq 2N_{ancestral}s < 1$ (i.e., effectively neutral mutations), $f_1$, with $1 \leq 2N_{ancestral}s < 10$ (i.e., weakly deleterious mutations), $f_2$, with $10 \leq 2N_{ancestral}s < 100$ (i.e., moderately deleterious mutations), and $f_3$, with $100 \leq 2N_{ancestral}s < 2N_{ancestral}$ (i.e., strongly deleterious mutations), where $N_{ancestral}$ was the initial population size and $s$ was the reduction in fitness of the mutant homozygote relative to wild-type. Data point represent the mean value, whilst error bars represent the standard deviation. $\theta = 2N\mu$ and $\rho = 2Nr$, where $N$ is the ancestral population size ($N_{ancestral}$), $\mu$ is the mutation rate, and $r$ is the rate of recombination. Variable rates are drawn from a uniform distribution, such that the mean rate across each simulation replicate was equal to the fixed rate, to enable fair comparisons (see the Materials and Methods section for further details).

**S21**

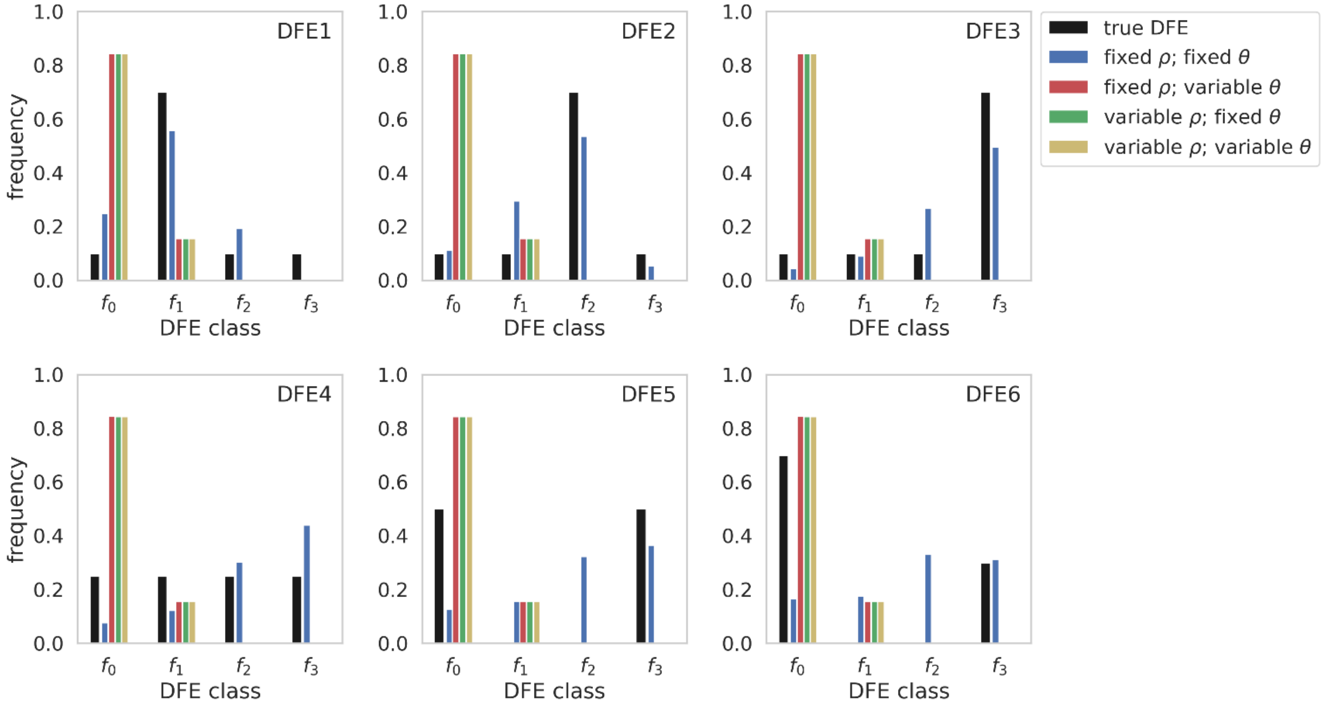


**S21:** Comparison of true distribution of fitness effects (DFEs) and DFE inference with **Grapes** when recombination and mutation rates are fixed and/or variable, under our six DFE models **for equilibrium population simulations**. Exonic mutations were drawn from a DFE comprised of four fixed classes (Johri et al. 2020), whose frequencies were denoted by $f_i$: $f_0$, with $0 \leq 2N_{ancestral}s < 1$ (i.e., effectively neutral mutations), $f_1$, with $1 \leq 2N_{ancestral}s < 10$ (i.e., weakly deleterious mutations), $f_2$, with $10 \leq 2N_{ancestral}s < 100$ (i.e., moderately deleterious mutations), and $f_3$, with $100 \leq 2N_{ancestral}s < 2N_{ancestral}$ (i.e., strongly deleterious mutations), where $N_{ancestral}$ was the initial population size and $s$ was the reduction in fitness of the mutant homozygote relative to wild-type. $\theta = 2N\mu$ *and* $\rho = 2Nr$, where N is the ancestral population size *(*$N_{ancestral}$*)*, $\mu$ is the mutation rate, and $r$ is the rate of recombination. Variable rates are drawn from a uniform distribution, such that the mean rate across each simulation replicate was equal to the fixed rate, to enable fair comparisons (see the Materials and Methods section for further details).

**S22**

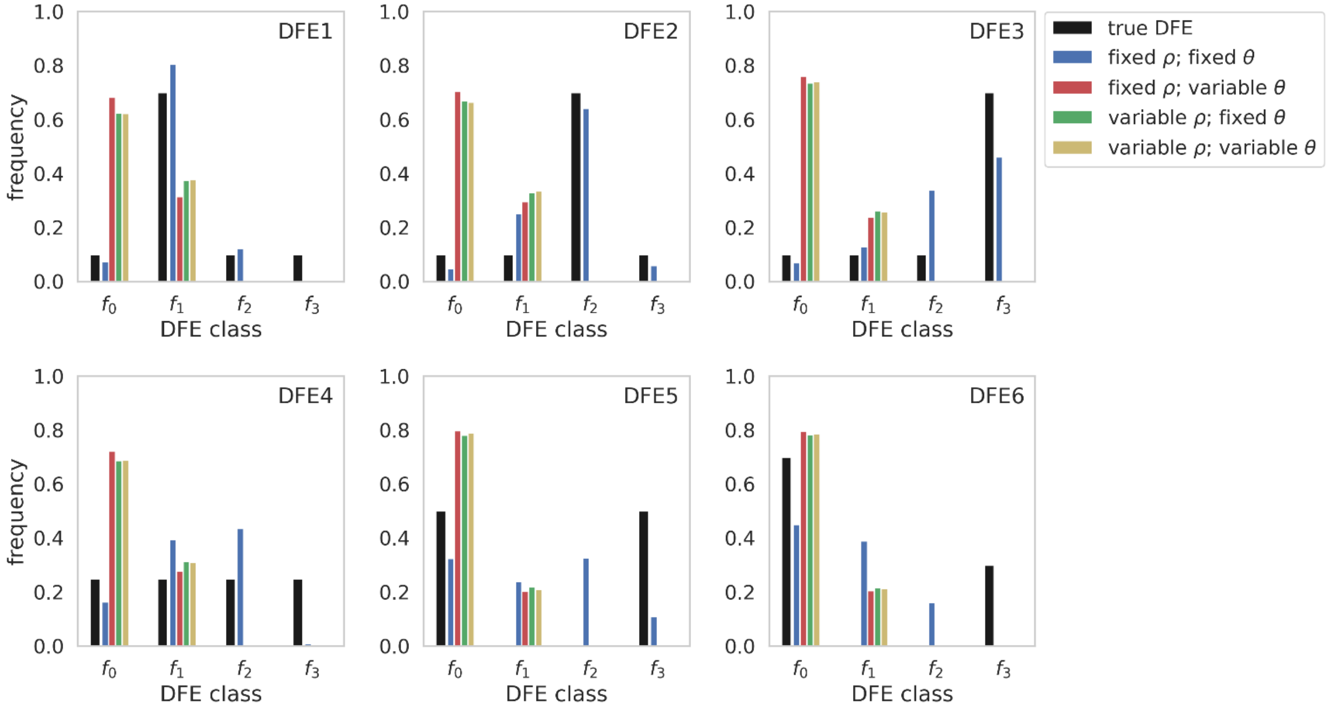


**S22:** Comparison of true distribution of fitness effects (DFEs) and DFE inference with **DFE-alpha** when recombination and mutation rates are fixed and/or variable, under our six DFE models **for population expansion simulations (such that $N_{current}$ = 2$N_{ancestral}$)**. Exonic mutations were drawn from a DFE comprised of four fixed classes (Johri et al. 2020), whose frequencies were denoted by $f_i$: $f_0$, with $0 \leq 2N_{ancestral}s < 1$ (i.e., effectively neutral mutations), $f_1$, with $1 \leq 2N_{ancestral}s < 10$ (i.e., weakly deleterious mutations), $f_2$, with $10 \leq 2N_{ancestral}s < 100$ (i.e., moderately deleterious mutations), and $f_3$, with $100 \leq 2N_{ancestral}s < 2N_{ancestral}$ (i.e., strongly deleterious mutations), where $N_{ancestral}$ was the initial population size and *s* was the reduction in fitness of the mutant homozygote relative to wild-type. $\theta = 2N\mu$ *and* $\rho = 2Nr$, where N is the ancestral population size *($N_{ancestral}$)*, $\mu$ is the mutation rate, and *r* is the rate of recombination. Variable rates are drawn from a uniform distribution, such that the mean rate across each simulation replicate was equal to the fixed rate, to enable fair comparisons (see the Materials and Methods section for further details).

**S23**

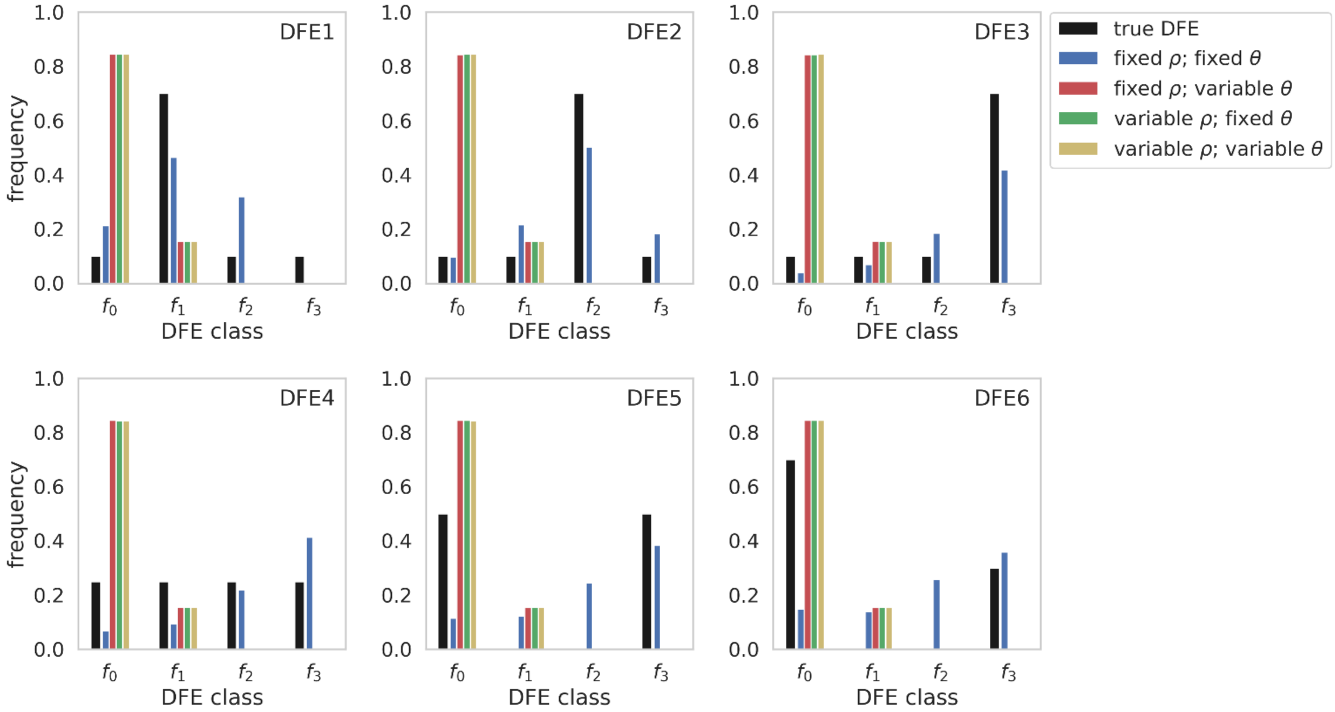


**S23:** Comparison of true distribution of fitness effects (DFEs) and DFE inference with **Grapes** when recombination and mutation rates are fixed and/or variable, under our six DFE models **for population expansion simulations (such that $N_{current}$ = 2$N_{ancestral}$)**. Exonic mutations were drawn from a DFE comprised of four fixed classes (Johri et al. 2020), whose frequencies were denoted by $f_i$: $f_0$, with $0 \leq 2N_{ancestral}s < 1$ (i.e., effectively neutral mutations), $f_1$, with $1 \leq 2N_{ancestral}s < 10$ (i.e., weakly deleterious mutations), $f_2$, with $10 \leq 2N_{ancestral}s < 100$ (i.e., moderately deleterious mutations), and $f_3$, with $100 \leq 2N_{ancestral}s < 2N_{ancestral}$ (i.e., strongly deleterious mutations), where $N_{ancestral}$ was the initial population size and $s$ was the reduction in fitness of the mutant homozygote relative to wild-type. $\theta = 2N\mu$ and $\rho = 2Nr$, where N is the ancestral population size ($N_{ancestral}$), $\mu$ is the mutation rate, and $r$ is the rate of recombination. Variable rates are drawn from a uniform distribution, such that the mean rate across each simulation replicate was equal to the fixed rate, to enable fair comparisons (see the Materials and Methods section for further details).

**S24**

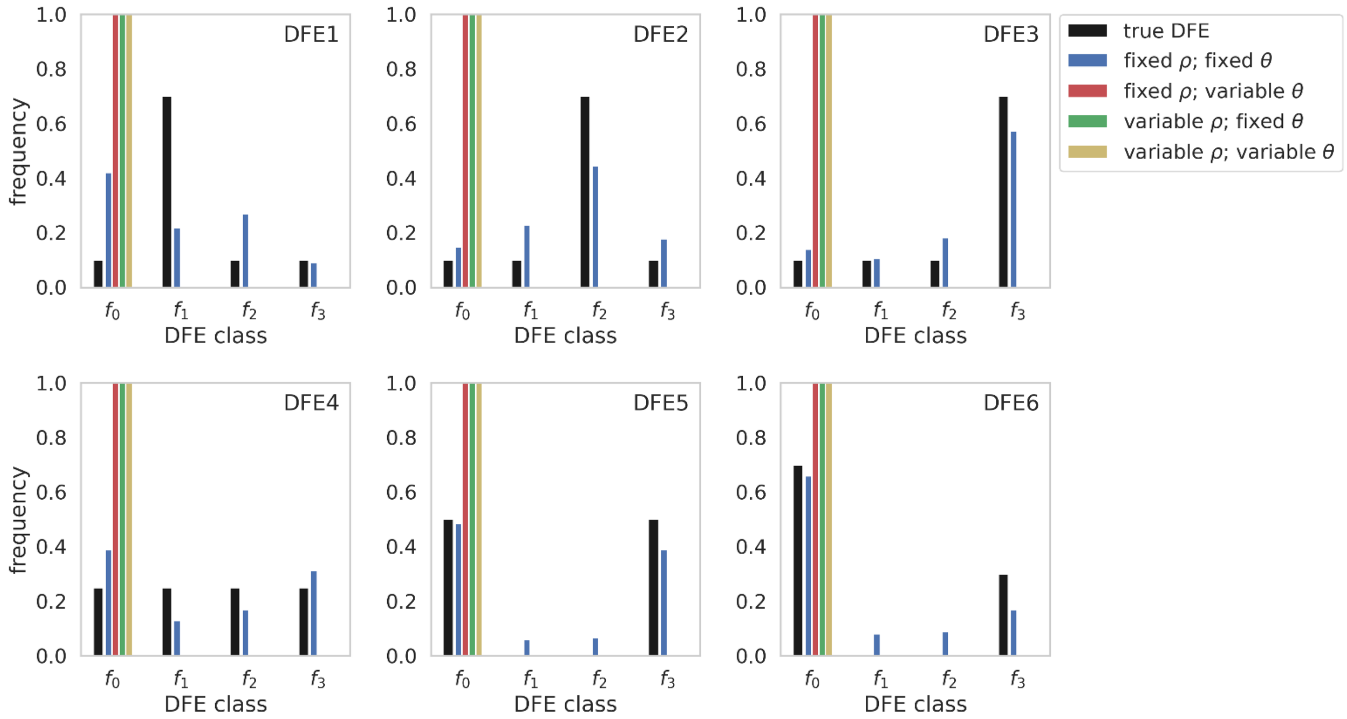


**S24:** Comparison of true distribution of fitness effects (DFEs) and DFE inference with **DFE-alpha** when recombination and mutation rates are fixed and/or variable, under our six DFE models **for population contraction simulations (such that $N_{current}$ = 0.5$N_{ancestral}$)**. Exonic mutations were drawn from a DFE comprised of four fixed classes (Johri et al. 2020), whose frequencies were denoted by $f_i$: $f_0$, with $0 \leq 2N_{ancestral}s < 1$ (i.e., effectively neutral mutations), $f_1$, with $1 \leq 2N_{ancestral}s < 10$ (i.e., weakly deleterious mutations), $f_2$, with $10 \leq 2N_{ancestral}s < 100$ (i.e., moderately deleterious mutations), and $f_3$, with $100 \leq 2N_{ancestral}s < 2N_{ancestral}$ (i.e., strongly deleterious mutations), where $N_{ancestral}$ was the initial population size and *s* was the reduction in fitness of the mutant homozygote relative to wild-type. $\theta = 2N\mu$ *and* $\rho = 2Nr$, where N is the ancestral population size *(*$N_{ancestral}$*)*, $\mu$ is the mutation rate, and *r* is the rate of recombination. Variable rates are drawn from a uniform distribution, such that the mean rate across each simulation replicate was equal to the fixed rate, to enable fair comparisons (see the Materials and Methods section for further details).

**S25**

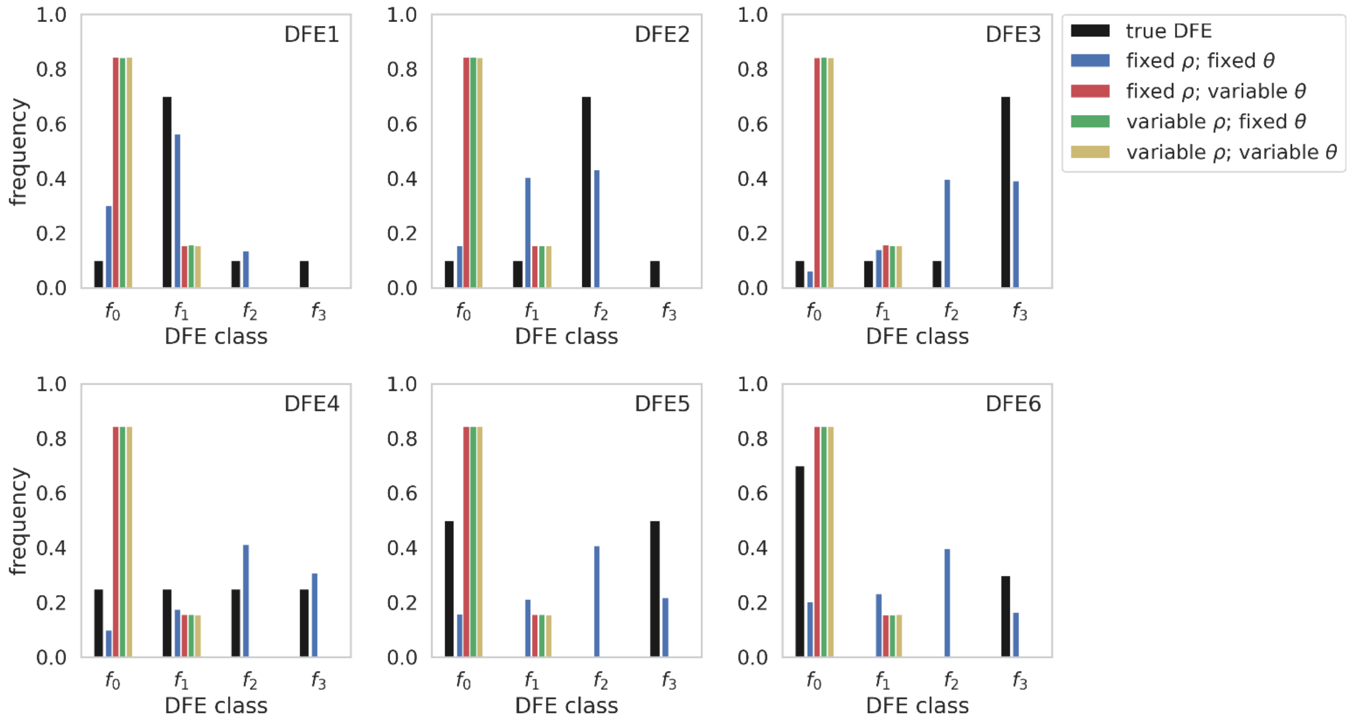


**S25:** Comparison of true distribution of fitness effects (DFEs) and DFE inference with **Grapes** when recombination and mutation rates are fixed and/or variable, under our six DFE models **for population contraction simulations (such that $N_{current}$ = 0.5$N_{ancestral}$).** Exonic mutations were drawn from a DFE comprised of four fixed classes (Johri et al. 2020), whose frequencies were denoted by $f_i$: $f_0$, with $0 \leq 2N_{ancestral}s < 1$ (i.e., effectively neutral mutations), $f_1$, with $1 \leq 2N_{ancestral}s < 10$ (i.e., weakly deleterious mutations), $f_2$, with $10 \leq 2N_{ancestral}s < 100$ (i.e., moderately deleterious mutations), and $f_3$, with $100 \leq 2N_{ancestral}s < 2N_{ancestral}$ (i.e., strongly deleterious mutations), where $N_{ancestral}$ was the initial population size and *s* was the reduction in fitness of the mutant homozygote relative to wild-type. $\theta = 2N\mu$ *and* $\rho = 2Nr$, where N is the ancestral population size *(*$N_{ancestral}$*)*, $\mu$ is the mutation rate, and *r* is the rate of recombination. Variable rates are drawn from a uniform distribution, such that the mean rate across each simulation replicate was equal to the fixed rate, to enable fair comparisons (see the Materials and Methods section for further details).

**S26**

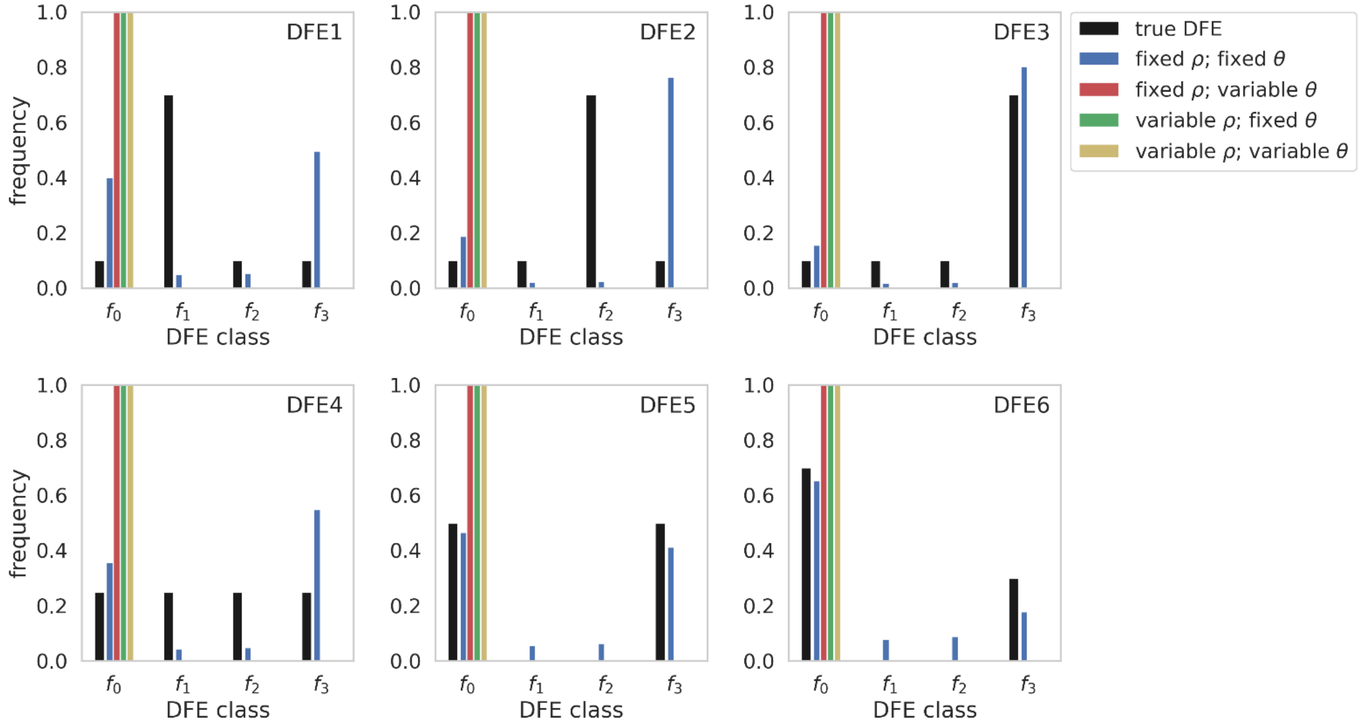


**S26:** Comparison of true distribution of fitness effects (DFEs) and DFE inference with **DFE-alpha** when recombination and mutation rates are fixed and/or variable, under our six DFE models **for population contraction simulations (such that $N_{current}$ = 0.01$N_{ancestral}$)**. Exonic mutations were drawn from a DFE comprised of four fixed classes (Johri et al. 2020), whose frequencies were denoted by $f_i$: $f_0$, with $0 \leq 2N_{ancestral}s < 1$ (i.e., effectively neutral mutations), $f_1$, with $1 \leq 2N_{ancestral}s < 10$ (i.e., weakly deleterious mutations), $f_2$, with $10 \leq 2N_{ancestral}s < 100$ (i.e., moderately deleterious mutations), and $f_3$, with $100 \leq 2N_{ancestral}s < 2N_{ancestral}$ (i.e., strongly deleterious mutations), where $N_{ancestral}$ was the initial population size and *s* was the reduction in fitness of the mutant homozygote relative to wild-type. $\theta = 2N\mu$ *and* $\rho = 2Nr$, where N is the ancestral population size *(*$N_{ancestral}$*)*, $\mu$ is the mutation rate, and *r* is the rate of recombination. Variable rates are drawn from a uniform distribution, such that the mean rate across each simulation replicate was equal to the fixed rate, to enable fair comparisons (see the Materials and Methods section for further details).

**S27**

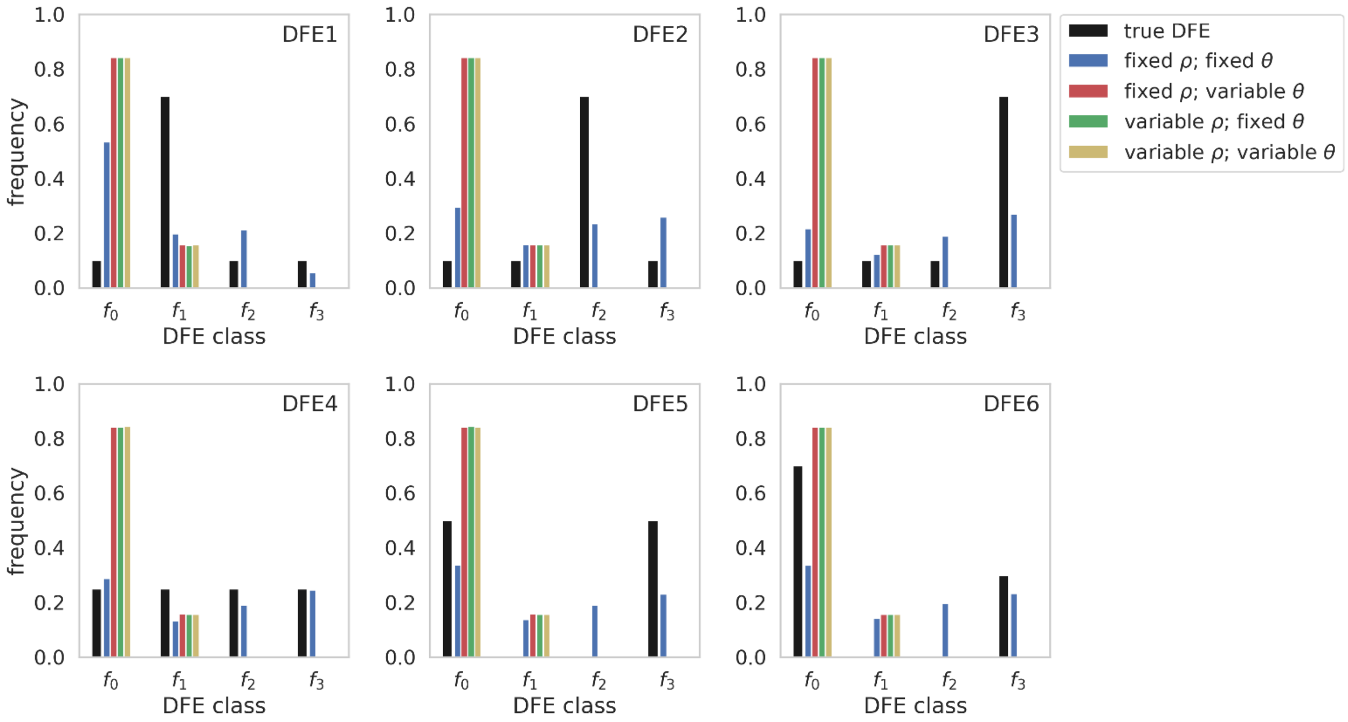


**S27:** Comparison of true distribution of fitness effects (DFEs) and DFE inference with **Grapes** when recombination and mutation rates are fixed and/or variable, under our six DFE models **for population contraction simulations (such that $N_{current}$ = 0.01$N_{ancestral}$)**. Exonic mutations were drawn from a DFE comprised of four fixed classes (Johri et al. 2020), whose frequencies were denoted by $f_i$: $f_0$, with $0 \le 2N_{ancestral}s < 1$ (i.e., effectively neutral mutations), $f_1$, with $1 \le 2N_{ancestral}s < 10$ (i.e., weakly deleterious mutations), $f_2$, with $10 \le 2N_{ancestral}s < 100$ (i.e., moderately deleterious mutations), and $f_3$, with $100 \le 2N_{ancestral}s < 2N_{ancestral}$ (i.e., strongly deleterious mutations), where $N_{ancestral}$ was the initial population size and *s* was the reduction in fitness of the mutant homozygote relative to wild-type. $\theta = 2N\mu$ *and* $\rho = 2Nr$, where N is the ancestral population size *(*$N_{ancestral}$*)*, $\mu$ is the mutation rate, and *r* is the rate of recombination. Variable rates are drawn from a uniform distribution, such that the mean rate across each simulation replicate was equal to the fixed rate, to enable fair comparisons (see the Materials and Methods section for further details).

**S28**

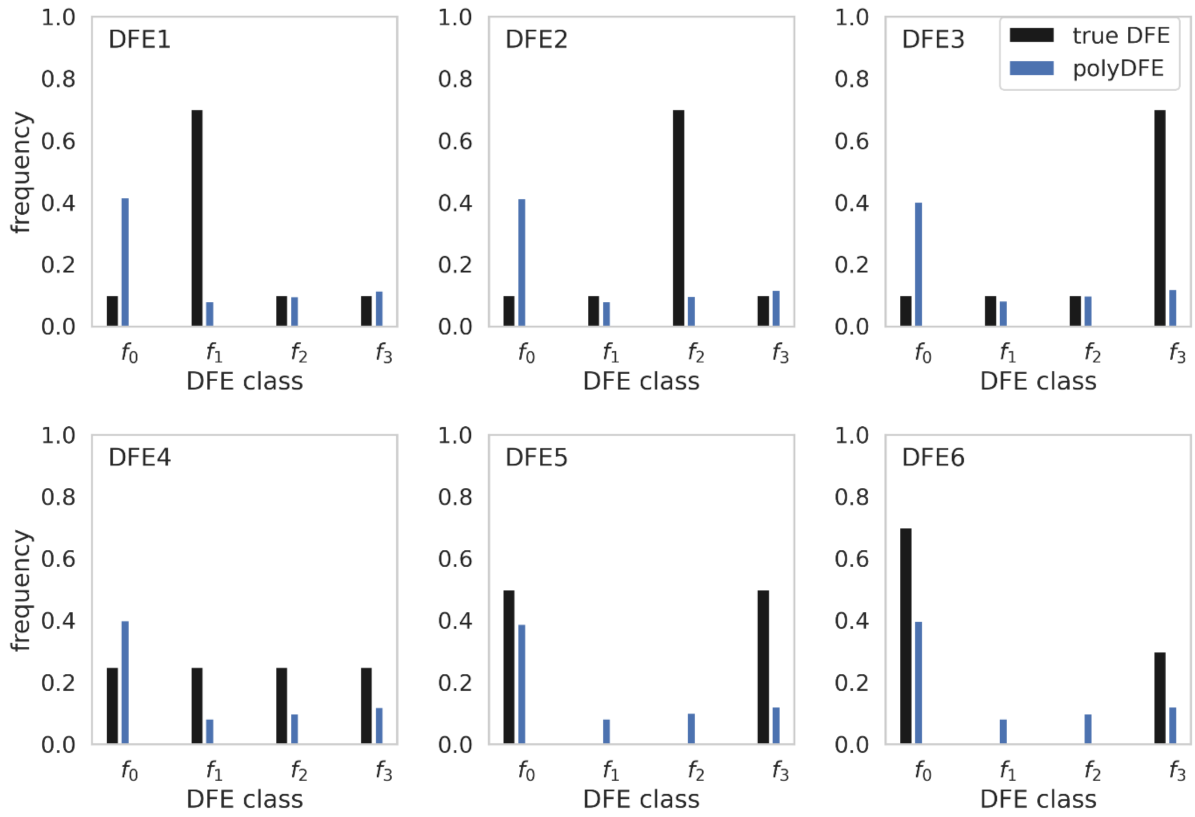


**S28:** Comparison of true distribution of fitness effects (DFEs) and DFE inference with **polyDFE** when recombination rates are fixed and mutation rates are variable, under our six DFE models **for equilibrium population simulations**. Exonic mutations were drawn from a DFE comprised of four fixed classes (Johri et al. 2020), whose frequencies were denoted by $f_i$: $f_0$, with $0 \leq 2N_{ancestral}s < 1$ (i.e., effectively neutral mutations), $f_1$, with $1 \leq 2N_{ancestral}s < 10$ (i.e., weakly deleterious mutations), $f_2$, with $10 \leq 2N_{ancestral}s < 100$ (i.e., moderately deleterious mutations), and $f_3$, with $100 \leq 2N_{ancestral}s < 2N_{ancestral}$ (i.e., strongly deleterious mutations), where $N_{ancestral}$ was the initial population size and $s$ was the reduction in fitness of the mutant homozygote relative to wild-type. $\theta = 2N\mu$ *and* $\rho = 2Nr$, where N is the ancestral population size ($N_{ancestral}$), $\mu$ is the mutation rate, and $r$ is the rate of recombination. Variable mutation rates are drawn from a uniform distribution, such that the mean rate across each simulation replicate was equal to the fixed rate, to enable fair comparisons (see the Materials and Methods section for further details).
